# Kinesin-5 promotes microtubule nucleation and assembly by stabilizing a lattice-competent conformation of tubulin

**DOI:** 10.1101/520072

**Authors:** Geng-Yuan Chen, Ana B. Asenjo, Yalei Chen, Jake Mascaro, David F. J. Arginteanu, Hernando Sosa, William O. Hancock

**Affiliations:** Department of Biomedical Engineering and Bioengineering, Pennsylvania State University, University Park, PA, 16802, USA; Department of Physiology and Biophysics, Albert Einstein College of Medicine, Bronx, NY, 10461, USA; Center for Bioinformatics, Department of Public Health Sciences, Henry Ford Health System, Detroit, MI, USA

**Keywords:** Kinesin, Eg5, microtubule, microtubule dynamics, tubulin wedge inhibitors, microtubule-associated proteins, coupled-equilibria

## Abstract

Besides sliding apart antiparallel microtubules during spindle elongation, the mitotic kinesin-5, Eg5 promotes microtubule polymerization, emphasizing its importance in mitotic spindle length control. Here, we characterize the Eg5 microtubule polymerase mechanism by assessing motor-induced changes in the longitudinal and lateral tubulin-tubulin bonds that form the microtubule lattice. Isolated Eg5 motor domains promote microtubule nucleation, growth and stability. Eg5 binds preferentially to microtubules over free tubulin, and colchicine-like inhibitors that stabilize the bent conformation of tubulin allosterically inhibit Eg5 binding, consistent with a model in which Eg5 induces a curved-to-straight transition in tubulin. Domain swap experiments establish that the family-specific Loop11, which resides near the nucleotide-sensing Switch-II domain, is necessary and sufficient for the polymerase activity of Eg5. Thus, we propose a microtubule polymerase mechanism in which Eg5 at the plus-end promotes a curved-to-straight transition in tubulin that enhances lateral bond formation and thereby promotes microtubule growth and stability.

## Introduction

Faithful segregation of genetic material to daughter cells requires tight control of mitotic spindle size and architecture. Stochastic switching between microtubule growth and shrinkage enables assembly and dynamic remodeling of the mitotic spindle, and these microtubule dynamics are mediated by dozens of regulatory proteins. Understanding the spatiotemporal regulation of microtubule dynamics during mitosis is essential for uncovering mechanisms that cells employ to ensure proper chromosome segregation.

Tubulin subunits are held in the microtubule lattice by both longitudinal tubulin-tubulin contacts that stabilize protofilaments, and lateral contacts that join adjacent protofilaments. Incorporation of tubulin into the lattice is determined by the tubulin nucleotide state, with the textbook explanation being that GTP-tubulin adopts a straight conformation that readily incorporates into the lattice, whereas GDP-tubulin adopts a curved conformation. However, more recent work has shown that soluble tubulin with either GTP- or GDP-bound adopt a curved conformation (Ayaz et al., 2012; Rice et al., 2008), suggesting that growth at the microtubule plus-end requires the incoming tubulin to undergo a curved-to-straight transition before being incorporated into the lattice. Consistent with this, images of growing microtubule plus-ends often show curled protofilaments in which longitudinal contacts have formed, but lateral contacts that require the curved-to-straight transition have not yet formed (McIntosh et al., 2018). One prediction of this model is that biasing the curved-to-straight transition of tubulin by mutagenesis (Geyer et al., 2015; Ti et al., 2016), small-molecule inhibitors (Gigant et al., 2005; Ravelli et al., 2004), or other means should affect lateral bond stability and generally promote microtubule polymerization.

Kinesin motors are one class of proteins that modulate mitotic microtubule dynamics, with the best-studied examples being the microtubule-depolymerizing kinesin-8s and kinesin-13s (Helenius et al., 2006; Varga et al., 2006, 2009). There is both structural and biochemical evidence that these motors preferentially bind to free tubulin and stabilize the curved conformation that is incompatible with lateral tubulin-tubulin bond formation (Arellano-Santoyo et al., 2017; Benoit et al., 2018; Wang et al., 2016a). There is also evidence that other kinesins, such as the kinesin-7 CENP-E, which is essential for proper chromosome alignment, accumulate at microtubule plus-ends and enhance microtubule stability in vivo (Gudimchuk et al., 2013). However, the underlying microtubule stabilization mechanisms remain unclear, and whether other mitotic kinesins also modulate tubulin dynamics is not established.

Tetrameric kinesin-5 is best known for sliding antiparallel microtubules apart during spindle formation, but in addition to its mechanical activities, there is evidence that kinesin-5 motors also alter microtubule dynamics. Based on knockouts of Cin8 or Kip1 in budding yeast, Gardner and colleagues argued that yeast kinesin-5 motors act as microtubule depolymerases during anaphase (Gardner et al., 2008). However, other studies in budding yeast, and work in fission yeast did not find evidence of fungal kinesin-5 depolymerase activity (Fridman et al., 2013; Rincon et al., 2017). In contrast, a dimeric construct of *Xenopus* Eg5 was shown to act as a microtubule polymerase *in vitro* (Chen and Hancock, 2015), an activity that was attributed to the ability of the two heads to “staple” together consecutive tubulin in a protofilament (Chen et al., 2016). The apparent discrepancy between the *in vivo* yeast results and the *in vitro* vertebrate results remains unresolved.

In this study, we reveal the mechanism of the microtubule polymerase activity of vertebrate Eg5. Contrary to the expectation that the two heads of Eg5 “staple” together adjacent tubulin dimers, we find that monomeric Eg5 motor domains strongly promote microtubule nucleation, increase the microtubule growth rate, and stabilize the microtubule lattice against depolymerization. Quantitative assays show that Eg5 monomers preferentially bind to straight rather than curved tubulin, consistent with the mechanism being a complement to the tubulin curvature-sensing mechanism proposed for the kinesin-8 and kinesin-13 microtubule depolymerases. We find that Loop11 is the key sequence in Eg5 that confers polymerase activity, and that swapping this structural element into kinesin-1 confers polymerase activity on this transport motor. Taken together, our results illustrate that Eg5 drives a curved-to-straight transition in tubulin at the plus-end of a growing microtubule, which strengthens lateral tubulin-tubulin contacts and promotes microtubule assembly. This microtubule polymerase activity may play an important role in the ability of kinesin-5 to control spindle size and architecture in mitotic cells.

## Results

### Eg5 motors enhance microtubule plus-tip stability

We previously showed dimeric *Xenopus* Eg5 accumulates at microtubule plus-ends, enhances the microtubule growth rate and decreases the microtubule catastrophe frequency (Chen and Hancock, 2015). A subsequent mechanochemical analysis indicated that the Eg5 predominantly resides in a two-heads-bound state (Chen et al., 2016), leading to the hypothesis that the polymerase activity results from the motor “stapling together” consecutive tubulin in a protofilament. To better understand the mechanism of Eg5 microtubule polymerization enhancement, we visualized microtubules grown in the presence and absence of dimeric Eg5 using both total-internal reflective fluorescence (TIRF) microscopy and negative-stain electron microscopy (EM). When microtubules were polymerized from unlabeled tubulin in the presence of GFP-labeled Eg5 dimers (Chen and Hancock, 2015), motors were observed streaming along the microtubules and 20% of growing plus-ends (N = 16 out of 83 microtubules) had curved tapers that were observable by TIRF microscopy (**Figure 1A**). These plus-end curls had diameters of ~1 μm, much larger than “ram’s horn” structures observed on depolymerizing microtubules, and many of the curved protofilament bundles subsequently straightened and were incorporated into the microtubule lattice (**Movie 1, 2**). In control experiments with kinesin-1, no plus-end curls were observed (N = 101 microtubules; not shown). When microtubules were examined in greater detail with negative-stain EM, sheets consisting of bundles of protofilaments were observed at many microtubule plus-ends. In the presence of Eg5, both the frequency of plus-end sheets and the length of the sheets were enhanced (**Figure 1B-D**). Together, these results suggest that Eg5 motors stabilize microtubule plus-tip tapers.

**Figure 1.**
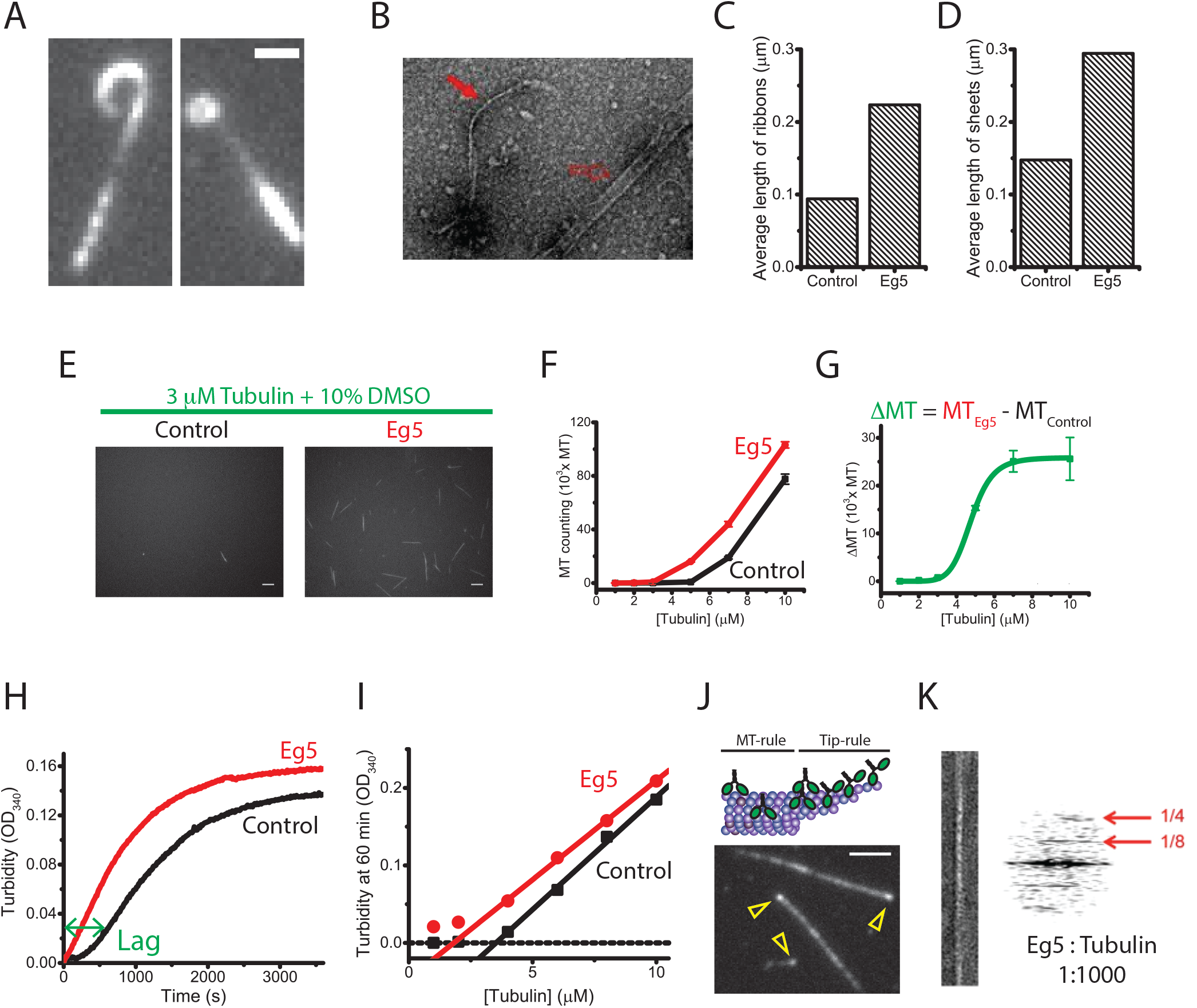
Eg5 motors enhance microtubule nucleation and growth. (A-D) Eg5 promotes diverse tip structures. (A) TIRF microscopy images of dynamic microtubule plus-ends growing in the presence of GFP-labeled dimeric Eg5. Scale bar: 1 *μ* m. (B) Negative-stain EM image of microtubules grown in the presence of Eg5, showing ribbon (closed arrow) and sheet (open arrow) structures. (C and D) Quantification of the mean length of ribbons and open sheets in the presence and absence of dimeric Eg5 motors (n_MT_ = 243-315). (E-I) Eg5 motors induce microtubule nucleation. (E) Epi-fluorescence images of TMR-labeled microtubules grown in the presence or absence of unlabeled Eg5 dimers (n = 10; mean ± SEM). Scale bar: 5 *μ* m. (F and G) Number of nucleated microtubules as a function of initial free tubulin concentration. (H) Averaged turbidity traces of 8 *μ* M free tubulin growing in the presence and absence of 130 nM Eg5 motors, demonstrating that Eg5 reduces nucleation lag time and results in higher turbidity plateau, indicating greater polymer mass (n = 3-5 traces). (I) Turbidity signal at 1 hour across tubulin concentrations demonstrating that Eg5 reduces critical concentration of tubulin nucleation. (J and K) Eg5 motors accumulate at the microtubule plus-ends. (J) Schematic and TIRF image of GFP-labeled Eg5 motors accumulating at the plus-end of a GMPCPP-stabilized microtubule. Open arrows indicate Eg5 end-bulbs. Scale bar: 2 *μ* m. (K) Negative-stain EM image and diffraction pattern of a straightened protofilament ribbon from microtubules grown at 1:1000 Eg5:tubulin stoichiometry. The strong 8 nm layer line observed is indicative of the motor decorating the protofilaments following the tubulin axial periodicity.

### Eg5 motors promote microtubule nucleation

Based on its stabilization of protofilaments at growing microtubule plus-ends, we hypothesized that Eg5 may also enhance microtubule nucleation (Roostalu and Surrey, 2017). To test this hypothesis, we polymerized microtubules starting from different free tubulin concentrations, immobilized the polymers on the surface of a coverslip, and counted the microtubules directly (Zheng et al., 1995). We found that at all tubulin concentrations, Eg5 motors increased the number of nucleated microtubules relative to control (**Figure 1E-G**). As an alternate method to assess nucleation, we carried out tubulin turbidity measurements to quantify the time-dependent increase in polymer mass, indicative of microtubule nucleation and growth (**Figure 1H**). In the absence of Eg5, the turbidity trace contains an initial lag (the nucleation phase), followed by a rising signal (the elongation phase) that eventually reaches a plateau. Addition of a physiologically relevant concentration of Eg5 in the reaction (130 nM) eliminated the lag phase, suggesting that Eg5 is indeed a microtubule nucleation enhancer (**Figure 1H**). Furthermore, the steady-state turbidity signals at 60 minutes increased across tubulin concentrations, indicating that Eg5 reduces the critical concentration of polymer formation (**Figure 1I**). The critical concentration in the presence of Eg5 was about half of the control (1.8 μM versus 3.5 μM), suggesting that Eg5 lowers the activation energy barrier of microtubule seed formation. Additionally, we measured microtubule growth from short GMPCPP microtubule seeds (Wieczorek et al., 2015) and found that Eg5 increased the fraction seeds that supported new growth across a range of tubulin concentrations (**Figure S1**). These three lines of evidence confirm that Eg5 is both a *de novo* and a template-based microtubule nucleation promoter. In addition to the established antiparallel microtubule sliding activity of Eg5, this nucleating activity may play a role in regulating aster formation during prometaphase (Helmke and Heald, 2014).

To better understand the microtubule stabilization activity of Eg5, we used negative stain electron microscopy to visualize motors on growing microtubules. Consistent with previous fluorescence work that showed plus-end accumulation of Eg5 (**Figure 1J**) (Chen and Hancock, 2015), we found that even at 1:1000 Eg5-to-tubulin stoichiometry, dimeric Eg5 decorated the protofilament ribbons coming from the microtubules (**Figure 1K**). Because microtubules have a cooperative structure stabilized by tubulin-tubulin longitudinal and lateral contacts, motor accumulation might synergistically stabilize the bundles and promote microtubule assembly, and the end-accumulating Eg5 motors might readily “staple” the newly engaged tubulin.

### Monomeric Eg5 is sufficient to promote microtubule assembly

Because of its dimeric structure, we hypothesized that Eg5 promotes microtubule stabilization and nucleation by “stapling” consecutive tubulin on a given protofilament into the lattice. To test whether this crosslinking is necessary for Eg5 polymerase activity, we deleted the coiled-coil region to generate a monomeric Eg5 construct, Eg5_M_, and used TIRF microscopy to quantify the microtubule growth rate in the presence of monomeric or dimeric Eg5 (**Figure 2A-C**). In the first experiment, we maintained a constant 20 μM tubulin concentration and varied motor concentrations (**Figure 2D**). Dimeric Eg5 motors increased the microtubule growth rate by an order of magnitude at this tubulin concentration, with half-maximal activity achieved at an EC_50_ ~ 2 nM. This EC_50_ is considerably tighter than the motor’s reported microtubule affinity of ~100 nM (Chen et al., 2016), which is likely due to concentration of motors at the plus-end, as seen in **Figure 1J**. Surprisingly, monomeric Eg5_M_ at high concentrations also potently promoted microtubule assembly, establishing that stapling of adjacent tubulin by dimeric motors is not necessary for the Eg5 polymerase activity. The EC_50_ of ~ 250 nM was close to its microtubule affinity (KM = 500 nM, **Figure 2G**), suggesting that maximum polymerase activity of the monomer requires nearly saturated occupancy of the lattice. Nevertheless, this result establishes that there are intrinsic properties within the Eg5 head domain that can modulate tubulin dynamics. We also found that, like Eg5 dimer, Eg5_M_ shortened the lag time in the turbidity assay and enhanced microtubule numbers (**Figure S2A-D**), demonstrating that Eg5 microtubule nucleation activity also operates by a mechanism other than the dimerized motor domains crosslinking adjacent tubulin in the lattice.

**Figure 2.**
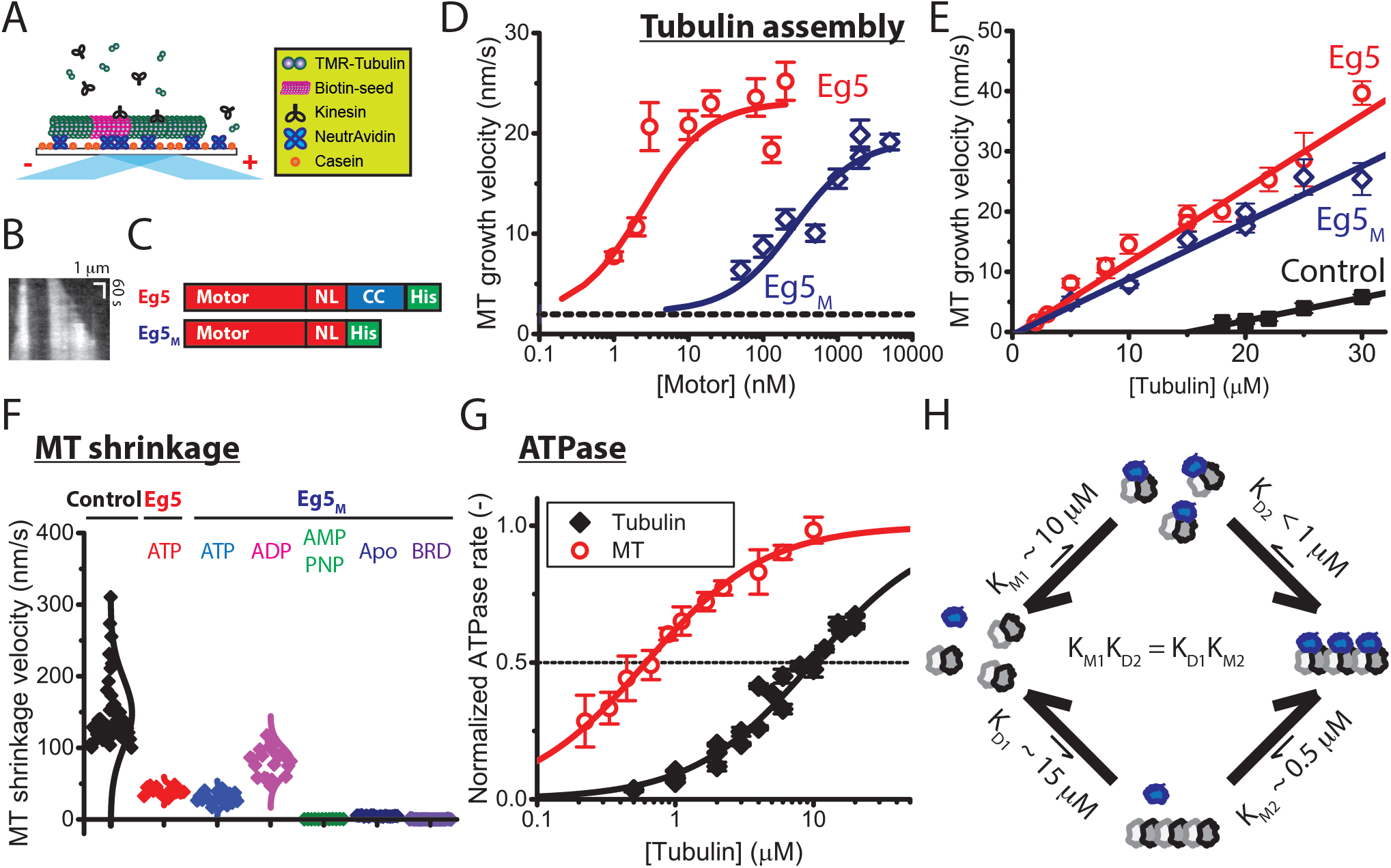
Monomeric Eg5 promotes microtubule assembly and enhances microtubule stability. (A-E) Microtubule growth rates. (A) Schematic of microtubule dynamics assays, in which TMR-labeled tubulin is grown off of biotinylated GMPCPP microtubule seeds and the microtubules are imaged by TIRF microscopy. (B) Example kymograph showing unlabeled seed and growth from both ends over time. (C) Diagrams of constructs. Dimeric Eg5 was made by replacing the native Eg5 coiled-coil with the kinesin-1 coiled-coil to ensure stable dimerization (Chen and Hancock, 2015; Shastry and Hancock, 2011). Monomeric Eg5_M_ was constructed by completely removing the Eg5 coiled-coil domain, leaving only the catalytic core and neck linker domains. (D) Motor-dependent microtubule growth velocities at 20 *μ* M tubulin and varying motor concentrations. EC_50_ values were 2.5 ± 0.6 nM for dimeric Eg5 and 260 ± 106 nM for monomeric Eg5 (mean ± SEM; n_MT_ = 6-51). (E) Tubulin-dependent microtubule elongation rates at saturating motor concentrations (130 nM Eg5 and 2 *μ* M Eg5_M_). From slopes of linear fits, apparent tubulin on-rates were 0.38 ± 0.04 *μ* M^−1^s^−1^, 1.22 ± 0.07 *μ* M^−1^s^−1^, and 0.93 ± 0.08 *μ* M^−1^s^−1^ for control, Eg5, and Eg5_M_, respectively. From y-intercepts, apparent off-rates were 5.82 ± 0.92 s^−1^, 0.65 ± 0.39 s^−1^, and 0.46 ± 1.16 s^−1^, and from x-intercepts, critical concentrations for tubulin assembly were calculated as 14 ± 1 *μ* M, 0.5 ± 0.3 *μ* M, and 0.5 ± 1.2 *μ* M for control, Eg5, and Eg5_M_, respectively (all mean ± SEM; n_MT_ = 6-40). (F) Effects of Eg5 on microtubule shrinkage rates following tubulin washout. Dimeric Eg5 was used at 80 nM and monomeric Eg5_M_ was used at 10 *μ* M under different nucleotide conditions (n_MT_ = 12-42). (G) Tubulin-and microtubule-stimulated ATPase of monomeric Eg5. ATPase rates are normalized to maximal for each substrate. Based on the K_M_ values, the apparent affinity of Eg5_M_ for tubulin (K_M1_ = 9.9 ± 1.2 μM) is 20-fold lower than its affinity for microtubules (K_M2_ = 0.6 ± 0.03 μM), which suggests that Eg5 motors favor the straight conformation of tubulin (mean ± SEM; n_MT_ = 5; n_Tub_ = 3). Mean ± SEM. (H) Schematic of affinity-driven polymer assembly. The twenty-fold difference of Eg5 affinity in **Figure 2G** generates a potential to facilitate tubulin assembly, consistent with the differential critical concentrations of microtubule growth in the presence and absence of monomeric Eg5 in **Figure 2E**.

To further explore effects of Eg5 on microtubule dynamics, we grew microtubules at varying tubulin concentrations in the presence and absence of saturating concentrations of motors (**Figure 2E**). Growth velocities are traditionally fit to a line, where the slope and y-intercept correspond to the tubulin on- and off-rate constants, respectively, and the x-intercept is the critical concentration for elongation. Under control conditions, the critical concentration for growth was 15 μM tubulin, but in the presence of high concentrations of either monomeric or dimeric Eg5, the critical concentration dropped to <1 μM tubulin, demonstrating that motor binding strongly drives microtubule growth. The apparent tubulin on-rate constant of 0.38 ± 0.04 μM^−1^s^−1^ in the absence of motors was increased to 1.22 ± 0.07 μM^−1^s^−1^ by dimeric Eg5 and to 0.93 ± 0.08 μM^−1^s^−1^ by Eg5_M_. More strikingly, the apparent tubulin off-rate of 5.82 ± 0.92 s^−1^ in the absence of motors was decreased ~10-fold to 0.65 ± 0.39 s^−1^ by Eg5 and to 0.46 ± 1.16 s^−1^ by Eg5_M_. Hence, the nearly 30-fold reduction in the critical concentration for growth is achieved by a moderate increase in the tubulin on-rate and a large decrease in the tubulin off-rate, consistent with both monomeric and dimeric Eg5 stabilizing tubulin in the microtubule lattice.

Because microtubule dynamic instability is determined by the competition between growth and shrinkage, greater net polymer mass can be achieved by either enhancing growth or inhibiting shrinkage (or both). Thus, after demonstrating that both dimeric and monomeric Eg5 promote microtubule growth, we next assessed whether Eg5 binding to the microtubule lattice enhances lattice stability. To test lattice stabilization, we carried out microtubule shrinkage assays by growing microtubules from immobilized seeds, washing out the free tubulin to induce catastrophe, and measuring the depolymerization rate in the absence and presence of motors (**Figure 2F**). Under control conditions, the shrinkage rate was 149 ± 7 nm/s, demonstrating that GDP-tubulin in the microtubule lattice rapidly dissociates from protofilament ends. In the presence of saturating concentrations of dimeric or monomeric Eg5 motors and ATP, the microtubule shrinkage rate slowed nearly 4-fold to 40 ± 2 nm/s and 30 ± 2 nm/s for Eg5 and Eg5_M_, respectively. Thus, monomeric Eg5 motor domains bound to the lattice enhance microtubule stability. We next tested the nucleotide dependence of lattice stabilization by Eg5_M_. In ADP, which induces a weak-binding state, the shrinkage rate was 2-fold slower than control (80 ± 4 nm/s; titration curve in **Figure S2E**). In contrast, AMPPNP, no nucleotide (apo), or the Eg5 inhibitor BRD9876, all of which induce strongly-bound states of the motor (Chen et al., 2017), had “taxane-like” effects, generating superstable microtubules (AMPPNP: 0.5 ± 0.05 nm/s; Apo: 4.7 ± 0.18 nm/s; BRD: 0.2 ± 0.07 nm/s; **Figure 2F**). Hence, Eg5 motor domains bound to the lattice trap tubulin in the polymer form and reduce tubulin dissociation rates, and the degree of lattice stabilization scales with the strength of motor binding to the lattice.

### Eg5 promotes tubulin assembly by preferentially binding to tubulin polymer

For microtubule-depolymerizing kinesins in the kinesin-8 and kinesin-13 families, the depolymerization mechanism can be explained by a thermodynamic cycle consisting of two linked equilibria, motor binding and microtubule assembly. In this formalism, the depolymerase activity can be explained as follows: motors bind more tightly to curved tubulin than to the microtubule lattice and thus motors shift tubulin toward the depolymerized state (Arellano-Santoyo et al., 2017; Benoit et al., 2018). By analogy, we propose a complementary mechanism for the Eg5_M_ polymerase activity – Eg5 motors bind more tightly to microtubule polymer than to free tubulin, and thereby drive the system toward the polymerized state. In **Figure 2H**, we present a thermodynamic scheme describing motor binding and microtubule polymerization. Binding affinities of motor to free tubulin and to the microtubule lattice are described by equilibrium constants K_M1_ and K_M2_, respectively. Microtubule polymerization from free tubulin is described as an equilibrium constant corresponding to the critical concentration for microtubule growth. From the x-intercepts in **Figure 2E**, the equilibrium constant under control conditions is K_D1_ = 15 μM, and the corresponding value in the presence of motors is K_D2_ = 0.5 μM. Thus Eg5_M_ binding shifts the apparent affinity for tubulin incorporation into the microtubule by 30-fold. If these linked equilibria are treated as a thermodynamic cycle, then it follows that:

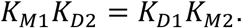

To quantify K_M1_ and K_M2_, the effective equilibrium constants for motor binding to tubulin and microtubules, respectively, we carried out ATPase assays and used the K_M_ as a proxy for the microtubule and tubulin affinity. From Eg5_M_ ATPase assays in **Figure 2G**, K_M1_ = 9.9 ± 1.2 μM for the tubulin-stimulated ATPase and K_M2_ = 0.6 ± 0.03 μM for the microtubule-stimulated ATPase. Thus, Eg5 motors prefer microtubules over tubulin as a substrate by nearly a factor of twenty. Plugging in the values, the products K_M1_ K_D2_ (9.9 μM * 0.5 μM) and K_D1_ K_M2_ (15 μM * 0.6 μM) agree within a factor of two. To summarize, preferential binding of Eg5 to microtubules over free tubulin (i.e. *K*_*M*2_ < *K*_*M*1_) enhances assembly of tubulin into the polymer state in the presence of Eg5 (i.e. *K*_*D*2_ < *K*_*D*1_). We next set out to understand the structural basis for the Eg5 microtubule polymerase mechanism.

### Eg5-binding causes a structural change in tubulin that promotes polymerization

In the microtubule lattice, tubulin heterodimers adopt a straight conformation stabilized by both lateral and longitudinal interactions with neighboring subunits, whereas tubulin in solution adopts a curved conformation incompatible with lateral contacts (Alushin et al., 2014; Ayaz et al., 2012; Rice et al., 2008). Thus, compounds that promote or stabilize the curved to straight transition generally promote polymerization, whereas compounds that stabilize the curved conformation inhibit polymerization (Akhmanova and Steinmetz, 2015). An example of the latter class of compounds are “intra-dimeric wedge inhibitors” such as colchicine derivatives and nocodazole that bind at the interface between α- and β-tubulin and are thought to freeze tubulin in the kinked conformation, which traps tubulins in the soluble state (Ravelli et al., 2004; Wang et al., 2016b).

Because monomeric Eg5 cannot directly crosslink adjacent tubulin-tubulin dimers, an alternate explanation is that Eg5 works allosterically by stabilizing the straight conformation of tubulin, which promotes lateral tubulin-tubulin interactions and thereby enhances microtubule growth. Structurally, the Eg5 binding site on the outer face of tubulin is located at the intradimeric junction between α- and β-tubulin, opposite to wedge inhibitor binding sites on the inner face of tubulin (**Figure 3A**). Based on this structural arrangement and the effects of different nucleotides on microtubule stabilization (**Figure 2F**), we put forward the following model. Eg5 binds free tubulin initially in a weakly-bound state, and upon tubulin binding, the motor releases its nucleotide and enters a strongly-bound state that stabilizes tubulin in a straight conformation and promotes microtubule assembly (**Figure 3B**). This model predicts that intradimeric wedge inhibitors like colchicine and nocodazole that stabilize the bent conformation of tubulin will antagonize with Eg5 for binding to tubulin and will inhibit tubulin-stimulated nucleotide release and ATP turnover.

**Figure 3.**
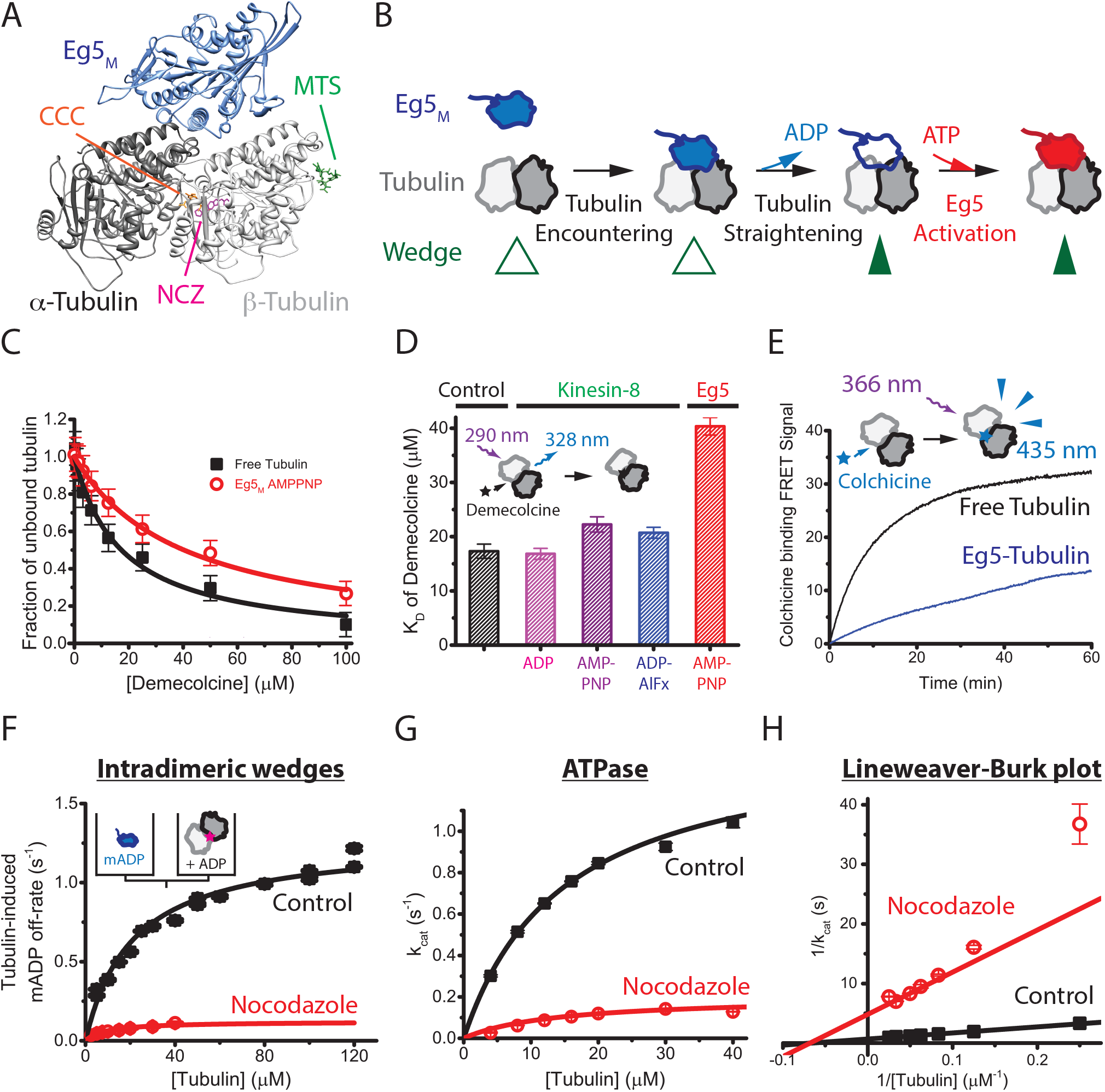
Eg5-binding causes a structural change of tubulin that promotes tubulin assembly. (A) Structure of Eg5 motor domain bound to free tubulin (PDB: 4AQW) and binding sites for inter- and intradimer wedge inhibitors. CCC, colchicine (PDB: 4O2B); NCZ, Nocodazole (PDB: 5CA1); MTS, Maytansinoid DM1 (PDB: 4TV8). (B) Model of coupling between nucleotide turnover in Eg5 and straightening of tubulin. Motor binding to free tubulin triggers ADP release, which causes a weak-to-strong binding transition of the motor that stabilizes the straight conformation of tubulin; ATP binding and hydrolysis follow. (C) Demecolcine binding assays based on quenching of tubulin autofluorescence by drug binding. Data were fit by a binding isotherm, yielding K_D_ of 17 ± 1.3 *μ* M for control and K_D_ of 40 ± 1.6 *μ* M in the presence of 4 *μ* M Eg5_M_ and 1 mM AMPPNP that induces the strong-binding state of the motor. (D) K_D_ for Demecolcine-tubulin binding, including results for the monomeric kinesin-8 KLP67A_M_ across various nucleotide-states; see **Figure S3A** for details. (E) Kinetics of colchicine binding to tubulin. 5 *μ* M tubulin was mixed with 50 *μ* M colchicine in the presence or absence of AMPPNP-bound motors, and the data fit by a rising biexponential. For control, fast and slow phases had rates and amplitudes of 0.169 ± 0.004 s^−1^ and 14 A.U., and 0.048 ± 0.001 s^−1^ and 19 A.U., respectively. In Eg5, fast and slow phases were 0.201 ± 0.022 s^−1^ and 1 A.U., and 0.0085 ± 0.0006 s^−1^ and 32 A.U., respectively. (F) Tubulin-induced mantADP release. Eg5 monomers pre-loaded with mantADP were flushed against tubulin pre-incubated with wedge inhibitors, and the resulting fluorescence fall due to mantADP release fit to a falling exponential. First-order rate constants where plotted across varying tubulin concentrations and fit to hyperbola to obtain the maximal ADP release rate, k_max_, and the tubulin concentration for half-maximal release, K_0.5_. Control: K_0.5_ = 21 ± 8 *μ* M, k_max_ = 1.3 ± 0.2 s^−1^; Nocodazole: K_0.5_ = 9 ± 13 *μ* M; k_max_ = 0.12 ± 0.06 s^−1^ (n = 5-7 for each averaged trace; mean ± SEM). (G and H) Tubulin-stimulated Eg5_M_ ATPase in the presence or absence of the intra-dimeric wedge inhibitor, nocodazole. Control: k_cat_ of 1.46 ± 0.04 s^−1^ and K_M_ of 14.9 ± 0.8 μM tubulin; nocodazole: k_cat_ of 0.20 ± 0.03 s^−1^ and K_M_ of 14 ± 5 μM tubulin; n = 3; mean ± SEM).

To test whether Eg5 binding induces a structural transition in tubulin, we first measured the binding affinity of the colchicine derivative demecolcine to tubulin in the presence or absence of motors (**Figure 3C**). We chose demecolcine because it binds tubulin with fast kinetics and it quenches tubulin intrinsic fluorescence, providing a convenient optical readout of drug binding (Pyles and Hastie, 1993). Demecolcine binds to free tubulin with a K_D_ of 17 ± 1.3 μM, and in the presence of the microtubule depolymerizing kinesin-8, KLP67A, the drug binding affinity was unaffected, consistent with drug and motor having distinct binding sites (**Figure 3D, S3A**). In contrast, in the presence of Eg5_M_ the K_D_ of demecolcine binding rose to 40 ± 1.6 μM, consistent with binding of Eg5 allosterically inhibiting drug binding (**Figure 3C, D**). Next, we measured the binding kinetics of colchicine to tubulin (**Figure 3E**). Previous work showed that the fluorescence signal consists of two phases, corresponding to an initial encounter, followed by a tubulin conformational change (Lambeir and Engelborghs, 1981). In the absence of Eg5, colchicine binding to tubulin was biexponential with a fast phase of 0.169 ± 0.004 s^−1^ and a slow phase of 0.048 ± 0.001 s^−1^ having equal amplitudes (**Figure 3E**). In the presence of Eg5, the amplitude of the fast phase dropped to ~3% of total, and the rate of the slow phase decreased fivefold to 0.0085 ± 0.0006 s^−1^. This result supports a model in which Eg5 binding straightens tubulin, closes the binding pocket of the wedge inhibitor, and generates a species that favors microtubule polymerization.

### Blocking the tubulin curved-to-straight transition slows the Eg5 nucleotide cycle

Because binding of Eg5 to tubulin diminishes the binding of tubulin wedge inhibitors, the converse should be true – tubulin wedge inhibitors should allosterically inhibit Eg5 binding to tubulin. Specifically, if Eg5-induced straightening of tubulin is coupled to the motor’s nucleotide hydrolysis cycle, then wedge inhibitors that stabilize the bent conformation of tubulin should diminish nucleotide release and ATP turnover by the motor. To measure motor-tubulin binding kinetics, we first incubated Eg5_M_ in the fluorescent nucleotide mantADP, and then flushed this species against varying concentrations of drug-bound tubulin plus unlabeled ADP. This experiment measures the kinetics of transitioning from State 1 to State 3 in **Figure 3B**. As a control experiment, we first tested the interdimeric wedge inhibitor maytansinoid DM1, which is thought to bind between tubulin dimers and inhibit polymerization by blocking longitudinal interactions rather than altering tubulin curvature. Control and maytansine kinetics were similar as expected (**Figure S3B**). In contrast, in the presence of the intradimeric wedge inhibitor nocodazole, which is thought to stabilize the curved conformation of tubulin, Eg5 nucleotide release was reduced more than fivefold (**Figure 3F**). This result is consistent with stabilization of the straight conformation of tubulin being coupled to nucleotide release by Eg5, which transitions the motor to the strong-binding state.

The coupling between tubulin straightening and Eg5 nucleotide turnover was also confirmed by solution ATPase assays. The tubulin-stimulated ATPase of Eg5_M_ was well-described by a Michaelis-Menten curve with K_M_ = 14.9 ± 0.8 μM and k_cat_ = 1.5 ± 0.04 s^−1^ (**Figure 3G, H**). The intradimeric wedge inhibitor nocodazole reduced the ATP turnover rate by 7-fold (K_M_ = 14.0 ± 5 μM, k_cat_ = 0.2 ± 0.03 s^−1^; **Figure 3H**). The identical x-intercepts (~−1/K_M_) in the Lineweaver-Burk plot indicate that nocodazole acts as a non-competitive inhibitor of the tubulin-stimulated Eg5_M_ ATPase (**Figure 3H**). The data are consistent with the Eg5 chemomechanical cycle being tightly coupled to a conformational change (possibly a curved-to-straight transition) in tubulin that is blocked by nocodazole binding.

### Loop11 mediates slow motility, end-binding, and microtubule polymerase activity of Eg5

In kinesins, Switch II in loop 11 plays a key role in mechanochemical coupling between the nucleotide binding and microtubule binding sites (Kull and Endow, 2002). Structurally, in tubulin-bound kinesin, Loop11 is located at the interface of alpha and beta tubulin (**Figure 4A**), placing it in the ideal location for detecting changes in tubulin curvature (Gigant et al., 2013). Notably, the microtubule depolymerizing kinesin-8, Kip3, has a strongly positively charged Loop11, and this loop has been shown to confer end-dwelling and microtubule depolymerase activities for this motor (Arellano-Santoyo et al., 2017). In contrast, Loop11 in kinesin-1, which lacks these activities, contains less positive charge. Loop 11 of Eg5 has a net +3 charge relative to kinesin-1, suggesting it has a greater electrostatic interaction with tubulin (**Figure 4A**). Hence, we tested the hypothesis that Loop 11 is a distinguishing feature of Eg5 that confers the motor’s microtubule polymerase activity.

**Figure 4.**
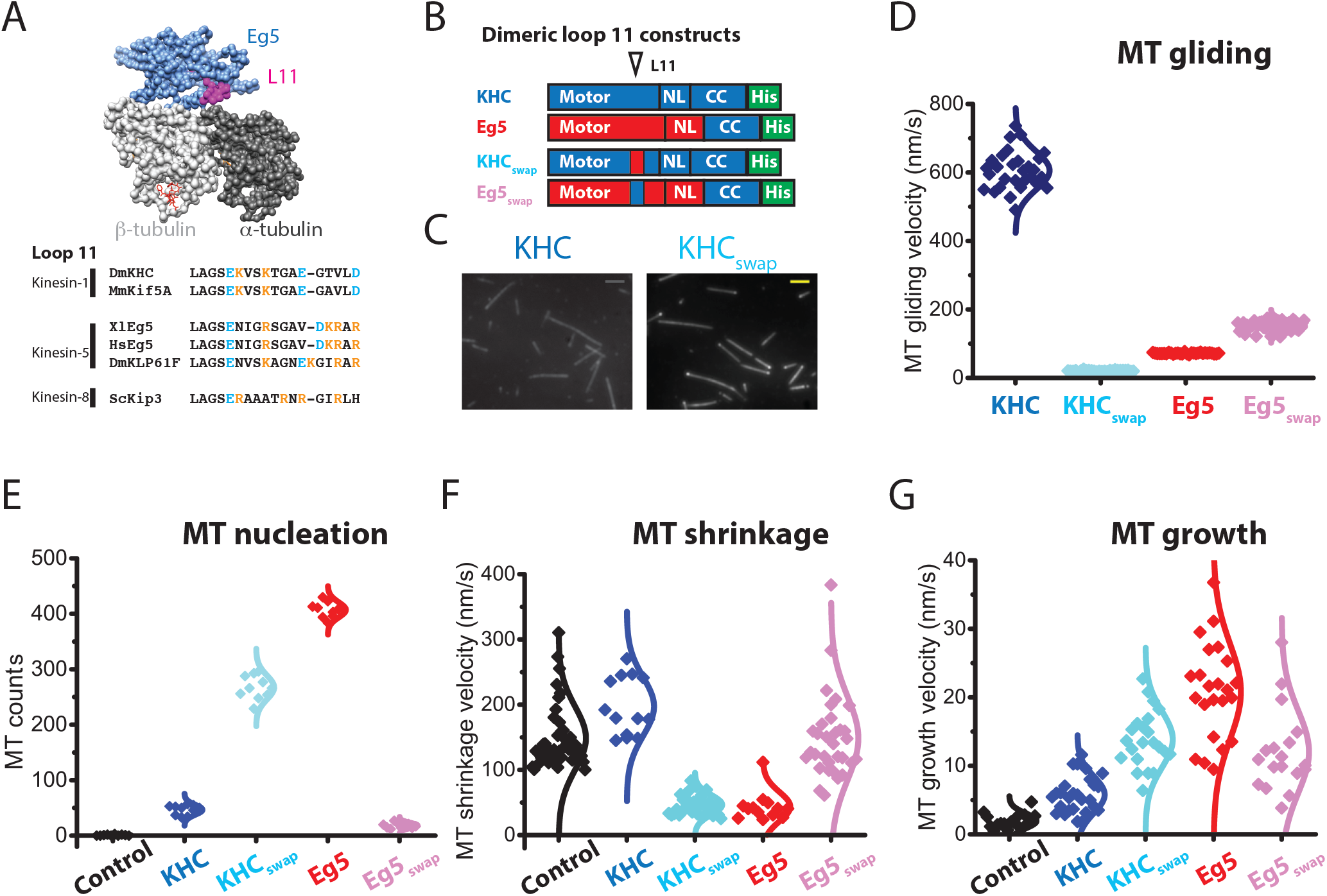
Loop11 mediates Eg5 end-accumulation and microtubule polymerase activity. (A) Structure of tubulin-bound Eg5, highlighting the location of Loop11 at the *α/β*-tubulin interface (PDB: 4AQW). Comparison of Loop11 sequences, highlighting positions of basic residues. Full alignments are shown in **Figure S4A**. (B) Diagram of Loop11-swapped mutants, KHC_swap_ and Eg5_swap_. (C) Swapping Eg5 Loop11 into KHC is sufficient to confer end-binding activity. Scale bar: 2 *μ* m. (D) Loop 11 strongly affects microtubule gliding speeds, consistent with it regulating the strong-binding state of the motor. Velocities: KHC, 606 ± 56 nm/s; KHC_swap_: 21 ± 2 nm/s; Eg5: 72 ± 2 nm/s; Eg5_swap_: 153 ± 16 nm/s; n_MT_ = 26-37. Mean ± SEM. (E) Number of newly formed microtubule across motor species. 5 *μ* M TMR-labeled microtubules plus 10 vol% DMSO were assembled in the presence or absence of 130 nM motors. Representative images are shown in **Figure S4B**. (F) Microtubule shrinkage rates following tubulin-washout, showing that Loop11 confers microtubule stabilization activity. Assays used 80 nM dimeric motors; control and Eg5 groups were taken from **Figure 2F** (n_MT_ = 13-42). (G) Microtubule growth rates showing that Loop11 contributes to growth enhancement activity of Eg5. Assays used 20 μM tubulin in the presence or absence of 10 nM dimeric motors; control and Eg5 groups were taken from **Figure 2D** (n_MT_ = 16-27).

To test the role of Loop 11, we swapped Loop 11 between the kinesin-1 KHC and Eg5 to create KHC_swap_ and Eg5_swap_ (**Figure 4B**). Our first observation was that, in contrast to wild-type KHC, KHC_swap_ accumulated at the plus-end of stabilized microtubules (**Figure 4C**; compare to Eg5 in **Figure 1J**). We next compared microtubule gliding velocities and found that, whereas wild-type KHC is 10-fold faster than Eg5, when their Loop 11 sequences were swapped, KHC was 7-fold slower than Eg5, indicating a profound change in chemomechanical coupling (**Figure 4D**). Previous work suggested that kinesin-1 spends half of its time in a one-head-bound state, whereas Eg5 spends the bulk of its stepping cycle in a two-heads-bound state (Andreasson et al., 2015; Chen et al., 2016; Mickolajczyk et al., 2015). Biochemical analysis (**Figure S4C-I**) indicates that KHC_swap_ spends the bulk of its cycle in a two-heads-bound state, whereas Eg5_swap_ becomes a one-head-bound motor. Additionally, differences in the Km values from ATPase assay demonstrates that KHC_swap_ favors straight lattices more than the curved tubulins (Tubulin: K_M_ = 48 ± 24 μM; Microtubule: K_M_ = 1.1 ± 0.2 μM; **Figure S4A**). Thus, Loop11 confers high microtubule affinity and end-dwelling activities.

We next tested the role of Loop11 on microtubule nucleation, growth and stabilization. Adding Eg5 to free tubulin led to significantly more nucleated microtubules than KHC, but the Loop11-swap mutants flipped this relationship (**Figure 4E**). Microtubule shrinkage following tubulin washout was slowed three-fold by Eg5 and not affected by KHC, whereas the relationship was flipped in the swap mutants (**Figure 4F**). Finally, Eg5 strongly enhanced and KHC only weakly enhanced the microtubule growth rate, and a swapping Loop11 flipped this relationship (**Figure 4G**). In summary, Loop11 in Eg5 is necessary for end-binding, microtubule stabilization, and enhancement of microtubule growth, and swapping in the Eg5 Loop11 is sufficient to confer these activities on the transport motor kinesin-1.

### Effect of Eg5 on MT dynamics can be explained by a 3 k_B_T enhancement of lateral tubulin bond energy

To gain quantitative insight into how Eg5 alters microtubule dynamics, we carried out numerical simulations of microtubule growth and shortening using a previously developed Monte Carlo model (Margolin et al., 2012). In this formalism, tubulin incorporation at a growing plus-end involves formation of an initial longitudinal bond in one protofilament that is subsequently stabilized by formation of a lateral bond with the adjacent protofilament (**Figure 5A**); shrinkage involves reversal of this pathway. Model parameters were taken from previous work or constrained by iterative fitting to our experimental data (**Table S1**). The model was able to recapitulate the experimental microtubule growth rates (**Figure 5B** and **C**) and shrinkage rates (**Figure 5D** and **Figure 5E**), both in the absence and presence of Eg5 motors. Remarkably, the observed effects of Eg5 on microtubule dynamics can be quantitatively explained by Eg5 binding causing a 10-fold enhancement of the tubulin curved-to-straight transition rate plus a 2.2-fold stabilization against lateral bond disruption. This is equivalent to Eg5 binding causing a 3.1 k_B_T enhancement of tubulin-tubulin lateral binding energy (**Table S2**). Hence, our results are quantitatively consistent with a ‘Zippering’ model in which Eg5 binding straightens tubulin, which promotes tubulin-tubulin lateral bond formation (**Figure 5F**). This lateral bond ‘zippering’ enhances the microtubule growth rate and stabilizes the microtubule lattice against depolymerization.

**Figure 5.**
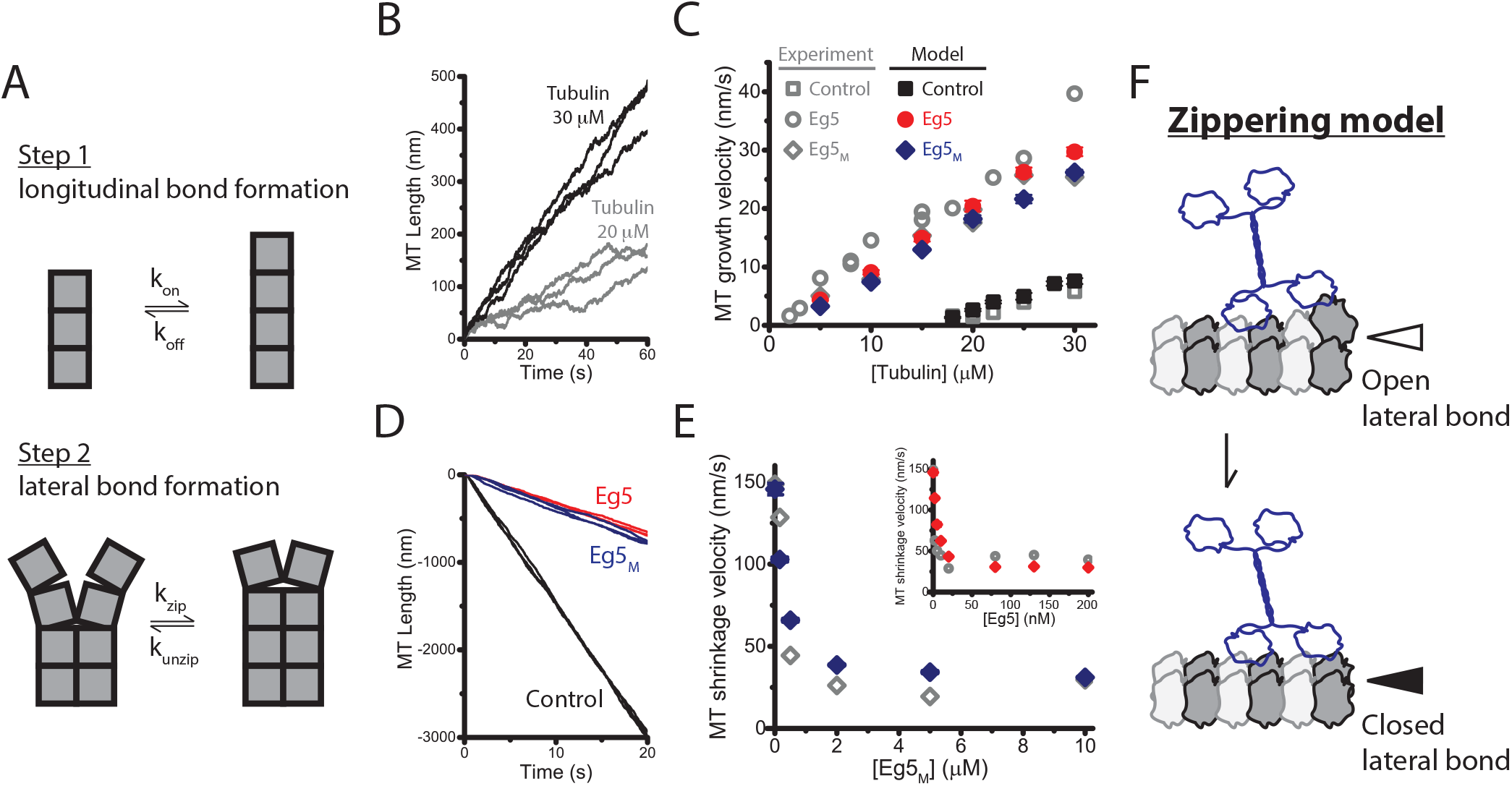
Eg5 polymerase can be explained by Eg5 enhancing tubulin lateral bond energy by 3 k_B_T. (A) Microtubule growth is simulated by separating tubulin incorporation at a growing tip into two steps. In Step 1, a single longitudinal bond is formed, with tubulin on- and off-rate constants *k_on_* and *k_off_*; and in Step 2 a lateral bond is formed with rate constants *k_zip_* and *k_unzip_*. The sequence is reversed during microtubule shrinkage. (B) Three simulated traces of microtubule growth at 20 *μ* M and 30 *μ* M tubulin from model in absence of Eg5. (C) Experimental (from **Figure 2E**) and simulated microtubule growth velocity across a range of tubulin concentrations. (D) Three simulated traces of microtubule shrinkage in the presence or absence of motors. (E) Experimental (from **Figure S2E**) and simulated shrinkage rates at varying Eg5_M_ (main figure) and Eg5 dimer (inset) concentrations. See **Figure** S5 for simulated kymographs and microtubule tip configurations. (F) Zippering model of Eg5-induced microtubule polymerization. Plus-end-bound Eg5 motors induce tubulin curved-to-straight transition at tip, thereby promoting lateral bond formation, which enhances the microtubule growth rate and stabilizes microtubules against shrinkage.

## Discussion

### A new role for Eg5 in mitotic spindle morphogenesis

A longstanding question in mitosis is how spindle size is regulated across cells of vastly varying dimensions. Two factors thought to be key to spindle size scaling are changes in the dynamic instability of the microtubules that make up the spindle, and rearrangement of the microtubules by antiparallel pushing forces generated by tetrameric kinesin-5 motors (Good et al., 2013; Helmke and Heald, 2014). In dividing cells, Eg5 localizes to the midzone during spindle formation, consistent with its mechanical role of sliding apart anti-parallel microtubules (Kapitein et al., 2005). Our finding that Eg5 enhances microtubule nucleation, polymerization and stability imply that the activities of this motor go beyond a simply mechanical role and suggest that Eg5 polymerase activity at the midzone may serve to maximize the degree microtubule overlap and drive poleward microtubule flux in the spindle (Miyamoto et al., 2004).

How do Eg5 motors promote microtubule assembly? Functional assays have shown that the microtubule detachment rate of Eg5 is relatively insensitive to force (Valentine et al., 2006), consistent with Eg5 motors primarily residing in a two-heads-bound “stapling” state that can resist load (Chen et al., 2016, 2017; Shimamoto et al., 2015). This “stapling” conformation implies a model in which the two heads cross-bridge consecutive tubulin dimers and thereby promote protofilament stability. Surprisingly, isolated Eg5 head domains, which bind to a single tubulin (Goulet et al., 2012), also promote microtubule polymerization, which rules out lattice crossbridging as the primary Eg5 polymerase mechanism. Hence, we considered a model in which Eg5 promotes microtubule polymerization by modulating the curvature of tubulin.

### Eg5 induces a curved-to-straight transition in tubulin that promotes polymerization

Because stability of the microtubule lattice result from the thermodynamics of the tubulin-tubulin contacts in the lattice (Gardner et al., 2011; Margolin et al., 2012), driving tubulin toward the straight conformation promotes microtubule assembly, whereas stabilizing the kinked conformation promotes disassembly. For instance, tubulin wedge inhibitors that stabilize the kinked conformation of tubulin, including colchicine, nocodazole, and vinblastine, disfavor lateral bond formation and thus decrease microtubule stability (Gigant et al., 2005; Ravelli et al., 2004; Wang et al., 2016b). Conversely, our finding that binding of Eg5 to tubulin reduces the binding affinity of tubulin wedge inhibitors suggests that motor binding induces a straightening of tubulin upon motor binding (**Figure 3C-D**).

In theory, Eg5 could induce a curved-to-straight transition of tubulin either by an induced-fit mechanism or by a conformational selection mechanism (Charles R. Cantor and Paul R. Schimmel, 1980). An induced-fit model suggests that Eg5 binding drives a curved-to-straight transition in tubulin, whereas a conformational selection model suggests that thermal fluctuations are sufficient to straighten tubulin and Eg5 motors selectively bind to and stabilize the straight conformation. Because structural studies suggest that tubulin wedge inhibitors only impact the bending flexibility but not the curvature (Ayaz et al., 2012; Gigant et al., 2005; Ravelli et al., 2004; Wang et al., 2016b), they can be used to distinguish between these alternate models, as follows. In **Figure 3F**, we used stopped-flow to measure the kinetics of Eg5-tubulin binding in the presence and absence of nocodazole, and found that Eg5 actually had a higher apparent affinity for nocodazole-bound tubulin than control tubulin. In a conformational selection model, decreasing tubulin flexibility would be predicted to reduce the probability of tubulin being in a straight conformation, and hence would predict a lower apparent motor binding affinity in the presence of drug. Thus, the higher apparent affinity is consistent with an induced fit model, and the finding that Eg5 nucleotide release is slower for nocodazole-bound tubulin suggests that Eg5 nucleotide release is coupled to the tubulin curved-to-straight transition induced by motor binding.

A model in which Eg5 binding induces a tubulin curved-to-straight transition is further substantiated by the long and flexible plus-end extensions seen on microtubules growing in the presence of Eg5 (**Figure 1A**). In published electron micrographs of isolated single protofilaments, they form curls and rings with diameters in the range of ~50 nm (McIntosh et al., 2018). The fact that plus-end curls observed here have diameters in the range of ~1 μm suggests that they are actually bundles of protofilaments. Recent high-resolution structural studies indicate that splayed protofilaments with “ram’s horn” geometries are present not only at the plus-ends of depolymerizing microtubules, but at the ends of assembling microtubules as well (McIntosh et al., 2018). These curved protofilaments suggest a two-step mechanism in which tubulin subunits are added to the end of growing protofilaments via longitudinal bonds, followed by a straightening step that results in lateral bond formation. In this scenario, Eg5 motors localized at the growing plus-end could bind to incoming tubulin and drive a curved-to-straight transition that would then accelerate lateral bond formation to generate protofilament bundles that anneal to form the intact microtubule lattice.

### Eg5 polymerase activity relies on a wedge-sensing structural element

Kinesin chemomechanical coupling relies on the concerted movement of the Switch-I and Switch-II domains that surround the nucleotide binding pocket (Nitta et al., 2008). Upon ATP hydrolysis, closure of Switch-I displaces Switch-II and results in a weak-binding motor. Because Switch-II dictates the kinesin strong binding state, it is reasonable to predict that family-specific Switch-II sequences might define microtubule polymerase activity of kinesin.

In structures of kinesin bound to tubulin, Loop11, which lies just proximal to Switch-II, is located at the interface of the alpha and beta subunits of tubulin, which is the fulcrum around which tubulin bending occurs (Arellano-Santoyo et al., 2017; Gigant et al., 2013). Sequence comparison across the kinesin superfamily showed that kinesin-5s, −7s, and −8s contain a strongly basic Loop11, whereas kinesin-1 transport motors do not (**Figure 4A, S4A**). We find that Loop11 is the key family-specific sequence in kinesin-5 that confers polymerase activity. This provides a complement to the microtubule depolymerizing kinesin-8, Kip3, where the family-specific Loop11 was required for that motor’s microtubule depolymerase activity (Arellano-Santoyo et al., 2017). Thus, Loop11 appears to be a key family-specific element that controls polymerase/depolymerase activity of kinesins. Interestingly, Loop11 sequences in both kinesin-5, −7s and kinesin-8 are considerably more basic than in kinesin-1, which suggests that tight electrostatic interactions between Loop11 and tubulin are essential for altering tubulin dynamics, but also that other subtle sequence differences define whether this tight binding results in stabilization or destabilization of the microtubule lattice.

### Multi-modal regulatory mechanisms of tubulin assembly

By considering motor binding and tubulin assembly as a coupled equilibrium, a general model emerges that can describe the mechanisms of both polymerizing and depolymerizing motors. We found that Eg5 motor domain preferentially binds to microtubules over tubulin, and that Eg5 enhances tubulin assembly. In the context of full-length tetrameric Eg5 in cells, the C-terminal motor tail, motor phosphorylation, as well as tubulin post-translational modifications or tubulin isoform differences may regulate the polymerase activity of Eg5. One general prediction is that any modification that preferentially enhances the affinity of the Eg5 motor domain for tubulin polymer should lead to greater polymerase activity. In contrast, the kinesin-8 Kip3, which depolymerizes microtubules was found to bind preferentially to free tubulin (Arellano-Santoyo et al., 2017). Similarly, depolymerizing kinesin-13 motors preferentially bind to tubulin curls over straight tubulin in the lattice (Friel and Howard, 2011). This thermodynamic model provides a general explanation for how other microtubule binding proteins may alter microtubule polymerization dynamics – those that bind more tightly to free tubulin promote depolymerization, whereas those that bind more tightly to the microtubule lattice promote polymerization. Furthermore, the model predicts that any modifications of Eg5 that alter the relative balance of tubulin- and microtubule affinity will alter the motor’s effects on microtubule polymerization.

In summary, our results support a model in which Eg5 binding to a free tubulin dimer or to a curved protofilament induces a curved-to-straight transition that promotes lateral assembly of protofilaments into a stable microtubule lattice. The effect of Eg5 on tubulin curvature relies on the Switch-II/Loop11 domains of Eg5 that are localized at the fulcrum between alpha and beta tubulin, a mechanism that is similar but complementary to the microtubule depolymerase mechanism of kinesin-8 motors. The effect of Eg5 on microtubule stability is reminiscent to the effects of taxane on microtubule stability, opening a possible strategy for anticancer therapeutics that alters microtubule stability through effects on the Eg5 motor.

## Acknowledgments

This work was supported by NIH R01 GM076476 and NIH R01 GM100076 to W.O.H. and R01 GM113164 to H.J.S. A portion of the bacterial culture work was carried out at the Penn State Huck Institutes for Life Sciences Fermentation Facility with the assistance of M. Signs. The authors thank Dan Sackett (NIH), David Sept (UMich), Luke Rice (UT Southwestern), Gary Brouhard (McGill), and John Correia (UMC) for thoughtful discussions on tubulin wedge inhibitors and tubulin curvature-sensing mechanisms.

## Author Contributions

G.-Y.C., Y.C., and W.O.H. initiated the project. G.-Y.C. and W.O.H. designed experiments, developed models, and prepared the manuscript. G.-Y.C. and D.F.J.A. performed protein purification and labeling, and G.-Y.C and J.M. carried out simulations and pilot studies of microtubule shrinkage. G.-Y.C. and A.B.A. generated samples for negative staining EM. A.B.A. and H.S. acquired and analyzed EM images.

## STAR Methods

### Materials and Methods

KEY RESOURCES TABLE

KEY RESOURCES TABLE
| REAGENT or RESOURCE | SOURCE | IDENTIFIER |
| --- | --- | --- |
| Antibodies |  |  |
| His-Tag Antibody | Thermo Fisher | R932-25 |
| Chemicals |  |  |
| GMPCPP | Jena Biosciences | NU-405L |
| GTP | Jena Biosciences | NU-1012-LYO |
| GDP | Jena Biosciences | NU-1172 |
| AMPPNP | Jena Biosciences | NU-214 |
| ATP | Sigma | A2383 |
| ADP | Sigma | A2754 |
| Mant-ADP | Thermo Fisher | M12416 |
| MgCl<sub>2</sub> | Quality Biological | 7058629 |
| DMSO | VWR | WN182 |
| BRD9876 | Tocris | 5454 |
| Colchicine | Sigma | C9754 |
| Demecolcine | Sigma | D7385 |
| Nocodazole | Sigma | M1404 |
| Maytansinoid DM1 | MedChemExpress | HY-19792 |
| Paclitaxel | Sigma | T7191 |
| α-casein | Sigma | C6780 |
| Catalase | Sigma | C1345 |
| Glucose Oxidase | Sigma | G6641 |
| 1H,1H,2H,2H-Perfluorodecyltrichlorosilane | Abcam | L16584 |
| β-mercaptoethanol | Sigma | M6250 |
| NeutrAvidin | Thermo Fisher | 31000 |
| Critical Commercial Assays |  |  |
| EnzChek | Thermo Fisher | E6646 |
| Q5 Site-directed Mutagenesis kit | NEB | E0554 |
| Experimental Models: Organisms/Strains |  |  |
| *E. coli* (BL21 DE3) | NEB |  |
| *E. coli* (5-α) | NEB | |
| Recombinant DNA |  |  |
| *Eg5*-GFP | Shastry and Hancock, 2011 |  |
| *KHC*-GFP | Shastry and Hancock, 2011 |  |
| *Eg5*-406 | Chen et al. 2015 |  |
| *KHC*-406 | Andreasson et al. 2015 |  |
| *Eg5<sub>M</sub>* | Chen et al. 2015 |  |
| *KHC<sub>M</sub>* | This study |  |
| *KLP67A<sub>M</sub>* | This study |  |
| *Eg5<sub>swap</sub>*-GFP | This study |  |
| *KHC<sub>swap</sub>*-GFP | This study |  |
| *Eg5<sub>swap</sub>* | This study |  |
| *KHC<sub>swap</sub>* | This study |  |
| Software and Algorithms |  |  |
| MATLAB | Mathworks | https://www.mathworks.com |
| ImageJ | NIH | https://imagej.nih.gov/ij/ |
| Micro-Manager | Vale Lab | https://micro-manager.org/ |
| MetaVue 6.3 | Molecular Devices | https://de.moleculardevices.com/ |
| Clustal Omega | EMBL-EBI | https://www.ebi.ac.uk/Tools/msa/clustalo/ |
| Soft MaxPro 5.4.6 | Molecular Devices | https://www.moleculardevices.com/ |
| Pro-Data SX/Viewer | Applied Photophysics | https://www.photophysics.com/ |
| RFPC 5301 | Shimadzu Scientific | https://www.ssi.shimadzu.com/support/ |
| OriginLab Pro 9.1 | OriginLab | https://www.originlab.com/ |
| UCSF Chimera | UCSF RBVI | https://www.cgl.ucsf.edu/chimera/ |

## CONTACT FOR REAGENT AND RESOURCE SHARING

Further information and requests for reagents may be directed to, and will be fulfilled by William O Hancock

## METHOD DETAILS

### Protein constructs, sequence design, and purification

Unlabeled KHC was generated by truncating *Drosophila* kinesin-1 dimer (KHC) to sequence 1-406 and fusing to a C-terminal hexaHis tag. GFP-labeled kinesin-1 (KHC-GFP) consisted of residues 1-560 of KHC linked to C-terminal eGFP and hexaHis tag. The monomeric kinesin-1 construct (KHC_M_) contains the head and neck linker domain (length 2-344) of KHC, preceded by an N-terminal hexaHis tag following the start codon. Eg5 dimers were generated by fusing the motor and neck linker domains (residues 1-368) of *Xenopus* kinesin-5 to the neck coil and coil 1 of KHC (residues 345-560), as previously described (Chen et al., 2016). Monomeric Eg5 (Eg5_M_) consisted of the head and neck linker (residues 1-368) of *Xenopus* kinesin-5. For Loop11 identification, kinesin-1, kinesin-5, and kinesin-8 sequences were compared using the Clustal Omega server (EMBL). To carry out Loop11-swapping, the primers with sequence of interest were synthesized (IDT) and introduced using the Q5 mutagenesis procedure (NEB). The kinesin-8 monomer (KLP67A_M_) consists of the head and neck linker of *Drosophila* KLP67A_1-360_. After PCR amplification, the linear DNAs were annealed using T4 ligase (NEB), followed by transformation into DH5α competent cells (NEB). All plasmids contain an ampicillin selection marker for antibiotic screening. All constructs were verified by sequencing (Penn State dnaTools).

Plasmids were transformed into E. coli BL21(DE3) for protein expression and grown in 2-liter cultures. Protein expression was induced by adding 1 mM IPTG and growing at 18 °C overnight. The collected pastes were resuspended into 25A200 buffer (25 mM K-ACES, pH 6.9, 2 mM Mg-Acetate, 2 mM K-EGTA, 0.1 mM K2-EDTA, 1 mM β-mercaptoethanol, 200 mM KCl, plus additional 10 μM ATP) and lysed by sonication. After Ni-column extraction, motors were exchanged into 0.5 μM mantADP with BRB80 buffer (80 mM PIPES, 1 mM EGTA, 1 mM MgCl_2_, pH 6.8) using GE HiTrap Desalting column, followed by adding 10 vol% sucrose as cryo-protectant for flash-freeze in liquid N_2_ before −80 °C storage. Detailed procedures for active motor quantification and yield optimization were described previously (Chen et al., 2015). All experiments were performed in BRB80 buffer at 23-24 °C unless otherwise noted.

PC-grade bovine brain tubulin was purified by three cycles of assembly and disassembly and labeled as described previously (Uppalapati et al., 2009). The labeled products were mixed with the unlabeled tubulin to generate aliquots of 5% Cy5-tubulin, 25% TMR-tubulin, and 5% biotinylated tubulin. GTP and Pi were removed by three additional assembly and disassembly rounds followed by buffer exchange into BRB80 plus 10 μM GDP. Tubulin concentrations were calculated by absorbance using an extinction coefficient ε_280_ = 115,000 M^−1^cm^−1^.

### Microtubule gliding assays

40 μM TMR-labeled tubulin, 8 vol% DMSO, and 2 mM Mg-GTP were incubated at 37°C and stabilized by adding 10 μM taxol. Motor gliding velocities of surface-adsorbed kinesin motors were imaged by TMR-labeled taxol-stabilized microtubules and quantified by MtrackJ plugin of ImageJ, as previously described (Chen et al., 2017). Unless otherwise indicated, the assays were carried out in imaging solution of BRB80 plus oxygen scavenger system and 0.2 mg/mL casein (Chen et al., 2017).

### Microtubule counting assays

TMR-labeled tubulin was incubated with 10 vol% DMSO plus 2 mM Mg-GTP in the presence or absence of kinesin motors plus 5 mM ATP at 37°C for 10 minutes to generate microtubule seeds. The nucleated products were diluted 10-fold in imaging solution plus 10 μM taxol at 23°C for 1 hour to elongate pre-formed seeds and minimize the *de novo* seed formation. The resulting solutions were diluted into imaging buffer plus 5 mM ATP and bound to the surface coated with 100 nM kinesin-1 rigor mutants for 20 minutes. The surface-bound microtubules with length >1 μm were scored.

### Microtubule dynamics and shrinkage assays

To form GMPCPP-seeds, 2 μM tubulin total (0.1 μM Cy5-labeled + 0.1 μM biotinylated + 1.8 μM unlabeled tubulin) were mixed with 0.2 mM Mg-GMPCPP at 37°C for 5 hours, followed by one spin-and-resuspension cycle to remove free tubulin. Coverslips were rinsed three times with 70% ethanol, followed by overnight acid cleaning in 6 M HCl. Acid-cleaned coverslips were rinsed by ddH_2_O, subjected to plasma cleaning (Harrick Plasma) for 2 minutes, incubated in vacuum-based dessicator with silane-vapor (Abcam) for at least 2 hours, and finally fixed by two pieces of double-sided tape to generate silanized flow chambers. To immobilize Cy5-labeled seeds, 500 nM neutravidin (Thermo Scientific), 5 wt% Pluronic F108 (BASF Corp.) 2 mg/mL casein were flowed in sequentially in intervals.

Polymerization was initiated by combining TMR-labeled tubulin, 5 mM Mg-ATP, and 2 mM Mg-GTP in the presence or absence of kinesin motors. For microtubule dynamics assays, the entire polymerization process was visualized under Nikon TE-2000 TIRF microscope, as previously described (Chen and Hancock, 2015). Microtubule growth velocities were measured by linear fit of kymographs using ImageJ software, where the fast growing ends were defined as the plus-ends.

For microtubule shrinkage assays, polymerization was carried out on a 37°C heat plate for 20 minutes, and catastrophe induced by flowing through room-temperature tubulin-free buffer. For experiments in presence of motors, motor concentrations in polymerization solution and shrinkage solution were identical to maintain motor-microtubule interactions both before and after catastrophe. The entire process was carried out on stage using epi-fluorescence microscope, as previously described (Uppalapati et al., 2009). Shrinkage velocities were quantified using the ImageJ plugin, MtrackJ.

### Kinesin tip-dwelling assays

Both Cy5-labeled GMPCPP-stabilized microtubules and Cy5-labeled taxol-stabilized microtubules were tested for motor tip-dwelling. To reach the steady-state end-bulbs, the GFP-labeled kinesin motors were incubated in the flow chamber for 10-15 minutes before image acquisition.

### Turbidity assays

Polymer formation was induced by mixing unlabeled tubulin plus 10 vol% DMSO, 2 mM Mg-GTP, 5 mM ATP, in the presence or absence of motors. Turbidity signals were recorded at 340 nm on a Multi-Mode Micro Plate Reader (Molecular Device Flexstation 3). The resulting traces were sigmoidal, corresponding to an initial nucleation phase followed by extension phase.

### Steady-state ATPase assays

Tubulin-stimulated ATPase rates were measured using the EnzChek phosphatase assay kit (Molecular probes, ThermoFisher Scientific). In solutions, 0.5 mM GDP and 3 mM ATP were used to avoid ATP-triggered tubulin polymerization. Microtubule-induced ATPase rates were carried out by enzyme-coupled assays with 5-20 nM active motors using a Multi-Mode Micro Plate Reader (Molecular Devices Flexstation 3), as previously described (Chen et al., 2017). Data are presented as mean ± s.e.m. for N = 3 determinations for each point, and fit with a Michaelis-Menten equation to obtain the maximal rate k_cat_ and the half-max concentration K_M_.

### Demecolcine-binding affinity and colchicine-binding kinetics

2 μM tubulin was first incubated with 3.5 μM kinesin motors plus nucleotides for 30 minutes, and then this tubulin-kinesin complex was mixed with varying concentrations of the colchicine analog, demecolcine, for the next 1 hour. Upon demecolcine binding, the quenching of tubulin intrinsic fluorescence at Ex 280 nm/ Em 346 nm provides a simple method to quantify binding fraction of demecolcine on tubulins. For colchicine-binding kinetics, 5 μM tubulin was first incubated with AMPPNP-bound Eg5 monomers for 30 minutes, followed by adding 50 μM colchicine and recording the tubulin-induced colchicine fluorescence signal (Ex 366 nm/ Em 435 nm). Traces were fit to a biexponential.

### Stopped flow experiments

Stopped-flow assays were performed on an Applied Physics SX20 spectrofluorometer, as previously described (Chen et al., 2015). Free tubulin was first incubated with 1.2x wedge inhibitors for 1 hour, and then flushed against 0.2 μM mantADP-bound Eg5 monomers. Upon tubulin binding, Eg5 motors release mantADP and result in a signal drop. Traces were fit to a mono-exponential to obtain the observed rate constants. Across varying tubulin concentrations, the observed rate constants follow the function,

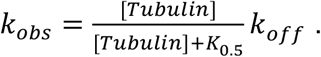

where *k_off_* and *K*_0.5_ denotes the ADP off-rate and the apparent MT affinity, respectively.

### Simulation of microtubule dynamics

Because tubulin has a kinked conformation in solution (Ayaz et al., 2012; Rice et al., 2008), two consecutive steps were included for tubulin incorporation into the lattice: longitudinal bond formation and lateral bond annealing (Margolin et al., 2012). Simulations started from a 13:2 seed that did not include a seam. The model required eight parameters to define microtubule dynamics: the tubulin-tubulin longitudinal on-rate (*k_on_*, μM^−1^s^−1^ per MT), the tubulin hydrolysis rate (*k_hyd_*, s^−1^), the longitudinal tubulin off-rates in GTP (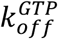, s^−1^) and GDP (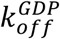, s^−1^), the forward and reverse lateral bond annealing rates (*k_zip_* and *k_unzip_*, s^−1^), and stabilization factors for both GTP tubulin (*F_GTP_*, fold) and the mechanical stabilization by the microtubule lattice (*F_wall_*, fold). To characterize microtubule shrinkage, the tubulin hydrolysis rate was set to either 50 s^-1^ or 0 s^−1^ to approximate GDP- or GMPCPP-tubulin, respectively; other parameters for growing microtubules are given in **Table S1**. To simulate microtubule growth and shortening in the presence of Eg5 motors, a further seven experimentally-derived (Chen and Hancock, 2015; Chen et al., 2016) or -constrained parameters were assigned: the Eg5 on-rate (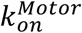, μM MT^−1^s^−1^), the Eg5 stepping rate on the lattice (*k_step_*, s^−1^) and on curved protofilaments (*k_slow_*, s^−1^), the Eg5 unbinding rate from the lattice (*k_unbind_*, s^−1^) and from curved protofilaments (*k_pause_*, s^−1^), a factor for the enhancement of lateral bond formation by Eg5 (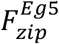, s^−1^), and a factor for the stabilization of lateral bonds by Eg5 (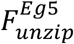, fold). The free energies resulting from these kinetic parameters are presented in **Table S2**.

### Structural superposition

Structures of Eg5-tubulin complex (PDB: 4AQW) and inhibitor-bound tubulins were aligned on the basis of a-tubulins (colchicine, PDB: 4O2B; nocodazole, PDB: 5CA1; Maytansinoid DM1, PDB: 4TV8) and visualized by UCSF Chimera.

### Quantification and statistical analysis

Data were fit in OriginPro 9.1 software. Unless otherwise indicated, data are presented as mean ± SEM.

## Supplemental Information Titles and Legends

**Figure S1. Related to Figure 1. Eg5 motors induce template-based nucleation**. (A) Epi-fluorescence images of TMR-labeled microtubules elongation on Cy5-labeled GMPPCP-seeds in the presence or absence of 130 nM unlabeled Eg5 dimers (n = 109-713; mean ± SEM). Scale bar: 5 μm. (B) Fraction of tip-nucleated microtubules as a function of initial free tubulin concentration.

**Figure S2. Related to Figure 2. Eg5 monomers promote polymer stability**. (A-D) Monomeric Eg5 promotes tubulin nucleation. (A) Averaged turbidity traces of 8 μM free tubulin polymerizing in the presence and absence of 2 μM Eg5 monomers, revealing that Eg5 promotes nucleation rate with a higher turbidity plateau (n = 3-5 traces). (B) Turbidity signal at 1 hour across tubulin concentrations demonstrating that monomeric Eg5 motors reduces critical concentration of tubulin polymerization. (C) Diagrams of constructs. Monomeric motors were generated by truncating kinesin coiled-coil domain and fused to a His-tag (Chen and Hancock, 2015). (D) Representative images and number of microtubules in the presence or absence of monomeric Eg5_M_ and KHC_M_. (E) Titration curve of monomeric Eg5 on dilution-induced microtubule shrinkage rates. The motor concentrations at half-max stability give a K_0.5_ of 1.7 ± 0.3 nM in 5 mM ADP and K_0.5_ = 0.3 ± 0.18 nM in 5 mM ATP. Inset: titration curve of dimeric Eg5 on dilution-induced microtubule shrinkage rates.

**Figure S3. Related to Figure 3. Eg5-modulated tubulin straightening perturbs colchicine-tubulin binding strength**. (A) Entire demecolcine binding isotherms shown in **Figure 3D**. (B) Tubulin-induced mantADP release in the presence and absence of Maytansine. K_0.5_. Control: K_0.5_ = 21 ± 8 μM, k_max_ = 1.3 ± 0.2 s^−1^; Maytansine: K_0.5_ = 20 ± 3 μM, k_max_ = 1.1 ± 0.07 s^−1^.

**Figure S4. Related to Figure 4. Loop11-mediated motor activity and microtubule stability**. (A) Sequence alignment of kinesin-1, −5, −7, and −8 motors. (B) Tubulin- and microtubule-stimulated ATPase of monomeric KHC_swap_, normalized to their maximal values. The K_M_ was 48 ± 24 μM on tubulins and 1.1 ± 0.2 μM on microtubules, suggesting that Eg5 motors favor microtubule lattices 40-fold over kinked, soluble tubulins (mean ± SEM; n_MT_ = 5; n_Tub_ = 3). (C) Representative images of microtubule number formed under varying motor species. (D-J) Loop11 regulates kinesin mechanical state transitioning. (D) Three-state mode includes motor one-head-bound and two-heads-bound transitions (Andreasson et al., 2015). The ATP-induced half-site release assays, in which mantADP-incubated one-head-bound motors were flushed against 2 mM ATP to release the tethered-head fluorescence signal, contain the transition time from the one-head-bound state to the two-heads-bound state (Chen et al., 2015; Hackney, 1994). The gliding velocity and its reciprocal represent the motor stepping rate and stepping time, respectively. The duration between the transient kinetics and the steady-state reaction time confers the reaction time of rear-head detachment. (E and F) Microtubule gliding assays of Loop11-swapped mutants (21 ± 2 nm/s for KHC_swap_ and 155 ± 15 nm/s for Eg5_swap_; n= 26-37, mean ± SD). The stepping times of KHC_swap_ and Eg5_swap_ were 381 ± 36 msec and 52 ± 5 msec, respectively. (G and H) ATP-induced half-site release assays of dimeric motors. The averaged traces were fit to a biexponential, where the rate of the fast phase corresponds to the first-passage time of the tethered-head touchdown (Chen et al., 2015). The ATP-induced motor binding rates of 140 ± 6 s^−1^ for KHC and 123 ± 11 s^−1^ for KHC_swap_ confer the tethered-head binding times of 7 ± 0.3 msec for KHC and 8 ± 0.7 msec for KHC_swap_, respectively. For the kinesin-5 motors, the rates of 24 ± 1 s^−1^ for Eg5 and 26 ± 3 s^−1^ for Eg5_swap_, meaning the reaction times of 42 ± 2 msec for Eg5 and 38 ± 4 msec for Eg5_swap_, indicate an unchanged transition time from the one-head-bound state to the two-heads-bound state. N = 5-7 for each determination. (I and J) The dominant state of Loop11-mutants. As listed in **Figure S4C**, KHC_swap_ motors spend ~98% in the two-heads-bound “staple” state, whereas Eg5_swap_ motors populate 73% in the one-head-bound state.

**Figure S5. Related to Figure 5. Parameterization of simulated microtubule dynamics**. (A) Simulated kymographs of growing and shrinking microtubules. (B) Illustration of Eg5-stimulated microtubule stability. Microtubule tip contains sheet structure in the presence of Eg5, whereas generates ram’s horn in control (Red and blue pair: GTP-tubulin; magenta and cyan pair: GDP-tubulin; green: two-heads-bound Eg5; black: one-head-bound Eg5). (C) EM images of shrinking microtubules in the presence or absence of monomeric Eg5. The diverse tips in the presence or absence of motors are consistent with the model illustration in **Figure S5C**. Scale bar: 100 nm.

**Table S1. Parameter set of Eg5-stimulated microtubule polymerization**.

**Table S2. Kinetics and free energy of tubulin lattice**

