## Supplemental Information for "Kinesin-5 promotes microtubule nucleation and assembly by stabilizing a lattice-competent conformation of tubulin"

### SUPPLEMENTARY RESULTS

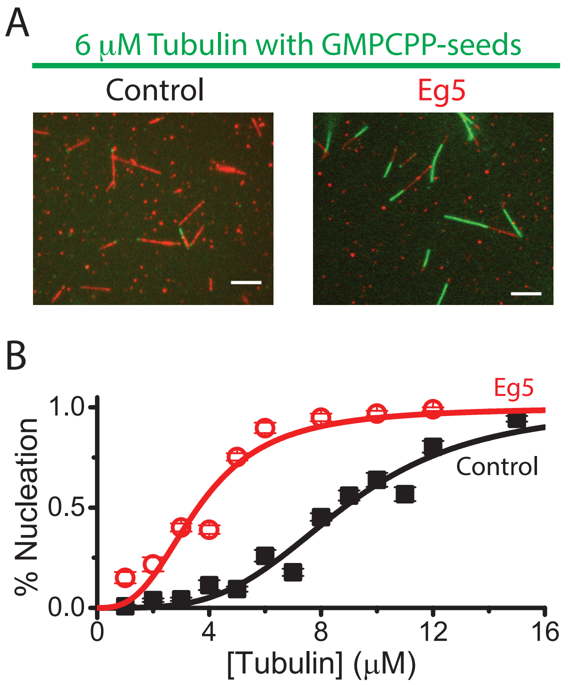


**Figure S1. Related to Figure 1. Eg5 motors induce template-based nucleation.** (A) Epi-fluorescence images of TMR-labeled microtubules elongation on Cy5-labeled GMPPCP-seeds in the presence or absence of 130 nM unlabeled Eg5 dimers (n = 109-713; mean ± SEM). Scale bar: 5 μm. (B) Fraction of tip-nucleated microtubules as a function of initial free tubulin concentration.

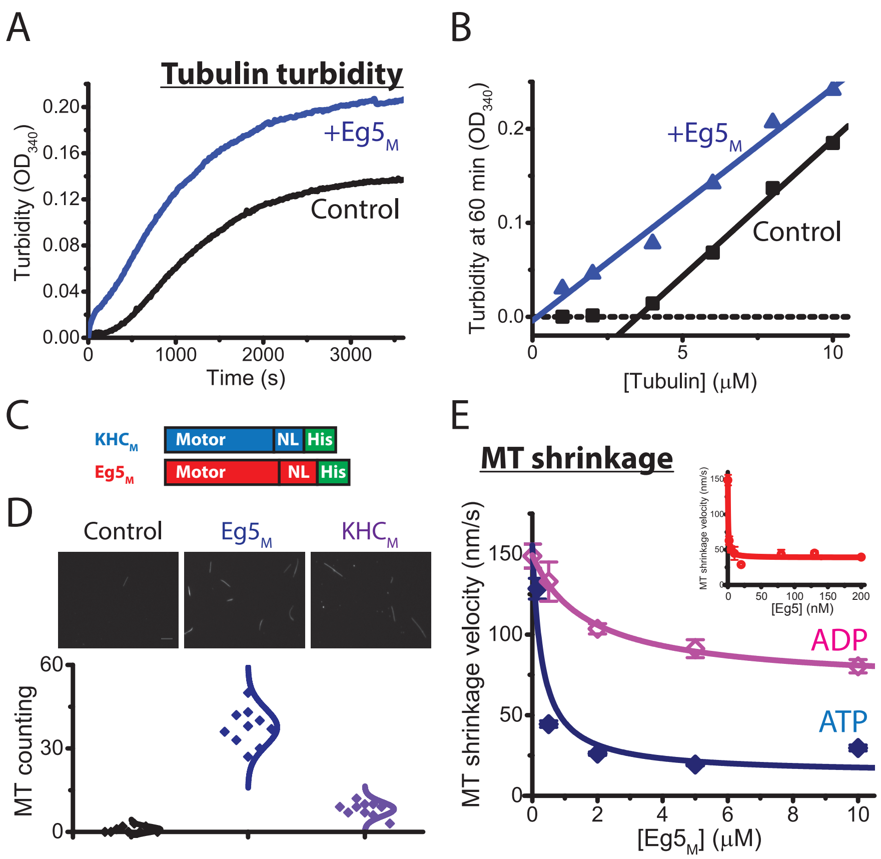


**Figure S2. Related to Figure 2. Eg5 monomers promote polymer stability.** (A-D) Monomeric Eg5 promotes tubulin nucleation. (A) Averaged turbidity traces of 8 μM free tubulin polymerizing in the presence and absence of 2 μM Eg5 monomers, revealing that Eg5 promotes nucleation rate with a higher turbidity plateau (n = 3-5 traces). (B) Turbidity signal at 1 hour across tubulin concentrations demonstrating that monomeric Eg5 motors reduces critical concentration of tubulin polymerization. (C) Diagrams of constructs. Monomeric motors were generated by truncating kinesin coiled-coil domain and fused to a His-tag (Chen and Hancock, 2015). (D) Representative images and number of microtubules in the presence or absence of monomeric $Eg5_M$ and $KHC_M$. (E) Titration curve of monomeric Eg5 on dilution-induced microtubule shrinkage rates. The motor concentrations at half-max stability give a $K_{0.5}$ of 1.7 ± 0.3 nM in 5 mM ADP and $K_{0.5}$ = 0.3 ± 0.18 nM in 5 mM ATP. Inset: titration curve of dimeric Eg5 on dilution-induced microtubule shrinkage rates.

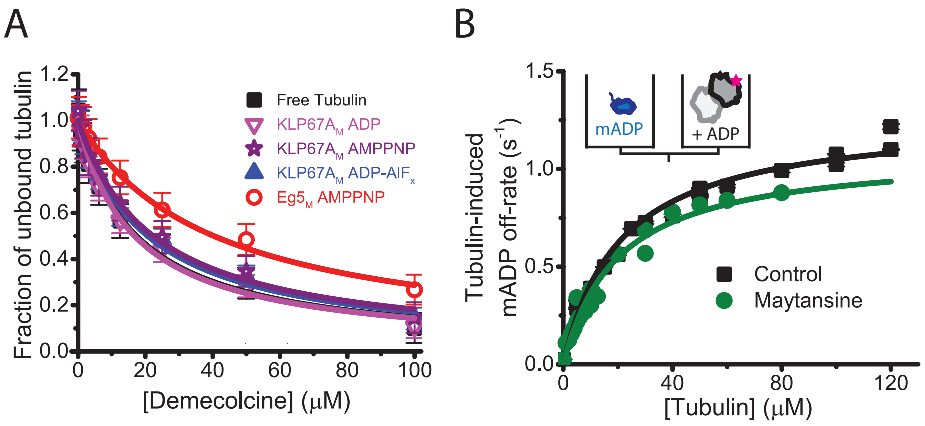


**Figure S3. Related to Figure 3. Eg5-modulated tubulin straightening perturbs colchicine-tubulin binding strength.** (A) Entire demecolcine binding isotherms shown in **Figure 3D**. (B) Tubulin-induced mantADP release in the presence and absence of Maytansine. $K_{0.5}$. Control: $K_{0.5}$ = 21 ± 8 μM, $k_{max}$ = 1.3 ± 0.2 $s^{-1}$; Maytansine: $K_{0.5}$ = 20 ± 3 μM, $k_{max}$ = 1.1 ± 0.07 $s^{-1}$.

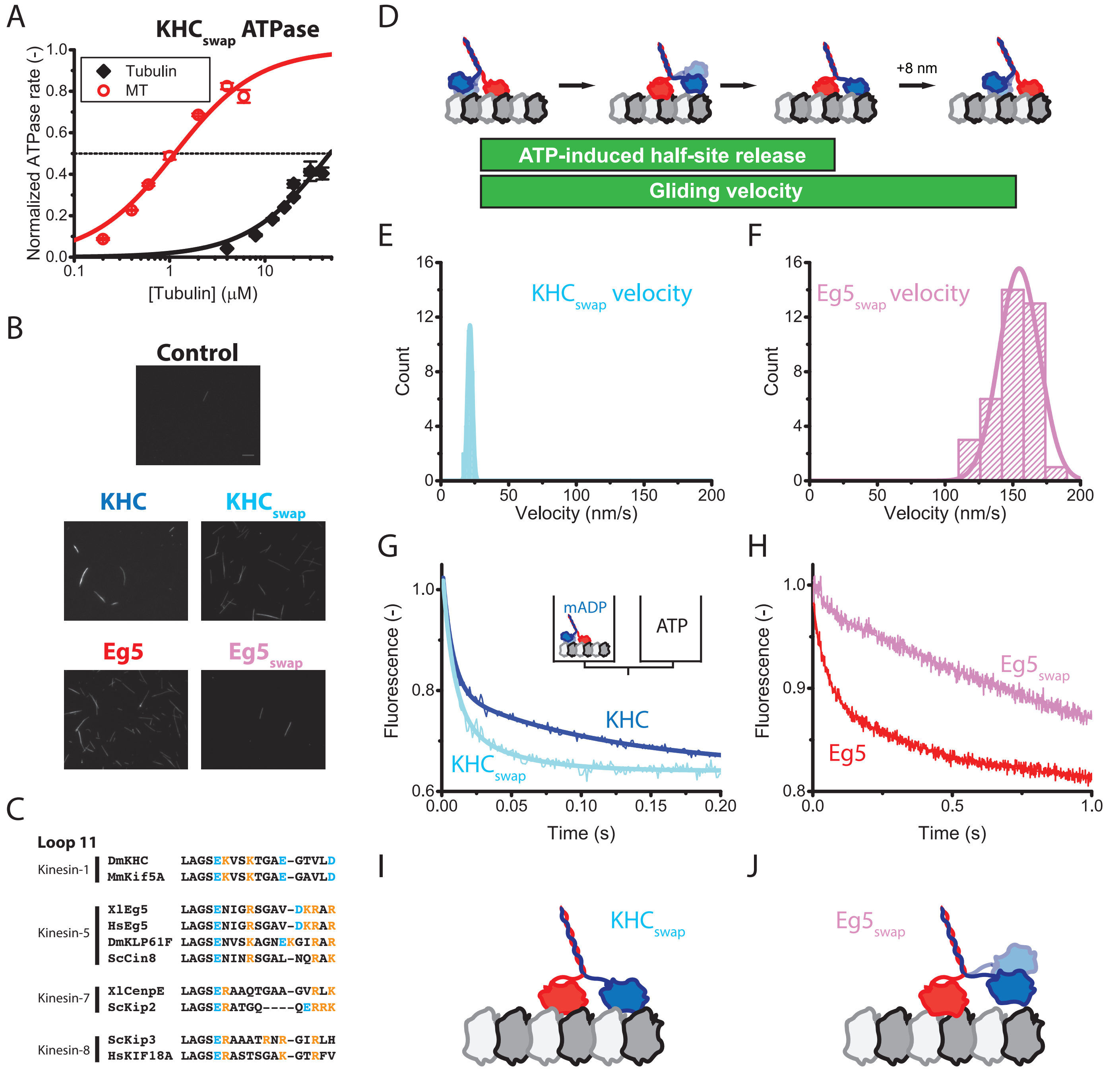


**Figure S4. Related to Figure 4. Loop11-mediated motor activity and microtubule stability.** (A) Sequence alignment of kinesin-1, -5, -7, and -8 motors. (B) Tubulin- and microtubule-stimulated ATPase of monomeric $KHC_{swap}$, normalized to their maximal values. The $K_M$ was 48 ± 24 μM on tubulins and 1.1 ± 0.2 μM on microtubules, suggesting that Eg5 motors favor microtubule lattices 40-fold over kinked, soluble tubulins (mean ± SEM; $n_{MT}$ = 5; $n_{Tub}$ = 3). (C) Representative images of microtubule number formed under varying motor species. (D-J) Loop11 regulates kinesin mechanical state transitioning. (D) Three-state mode includes motor one-head-bound and two-heads-bound transitions (Andreasson et al., 2015). The ATP-induced half-site release assays, in which mantADP-incubated one-head-bound motors were flushed against 2 mM ATP to release the tethered-head fluorescence signal, contain the transition time from the one-head-bound state to the two-heads-bound state (Chen et al., 2015; Hackney, 1994). The gliding velocity and its reciprocal represent the motor stepping rate and stepping time, respectively. The duration between the transient kinetics and the steady-state reaction time confers the reaction time of rear-head detachment. (E and F) Microtubule gliding assays of Loop11-swapped mutants (21 ± 2 nm/s for $KHC_{swap}$ and 155 ± 15 nm/s for $Eg5_{swap}$; n= 26-37, mean ± SD). The stepping times of $KHC_{swap}$ and $Eg5_{swap}$ were 381 ± 36 msec and 52 ± 5 msec, respectively. (G and H) ATP-induced half-site release assays of dimeric motors. The averaged traces were fit to a biexponential, where the rate of the fast phase corresponds to the first-passage time of the tethered-head touchdown (Chen et al., 2015). The ATP-induced motor binding rates of 140 ± 6 $s^{-1}$ for KHC and 123 ± 11 $s^{-1}$ for $KHC_{swap}$ confer the tethered-head binding times of 7 ± 0.3 msec for KHC and 8 ± 0.7 msec for $KHC_{swap}$, respectively. For the kinesin-5 motors, the rates of 24 ± 1 $s^{-1}$ for Eg5 and 26 ± 3 $s^{-1}$ for $Eg5_{swap}$, meaning the reaction times of 42 ± 2 msec for Eg5 and 38 ± 4 msec for $Eg5_{swap}$, indicate an unchanged transition time from the one-head-bound state to the two-heads-bound state. N = 5-7 for each determination. (I and J) The dominant state of Loop11-mutants. As listed in **Figure S4C**, $KHC_{swap}$ motors spend ~98% in the two-heads-bound "staple" state, whereas $Eg5_{swap}$ motors populate 73% in the one-head-bound state.

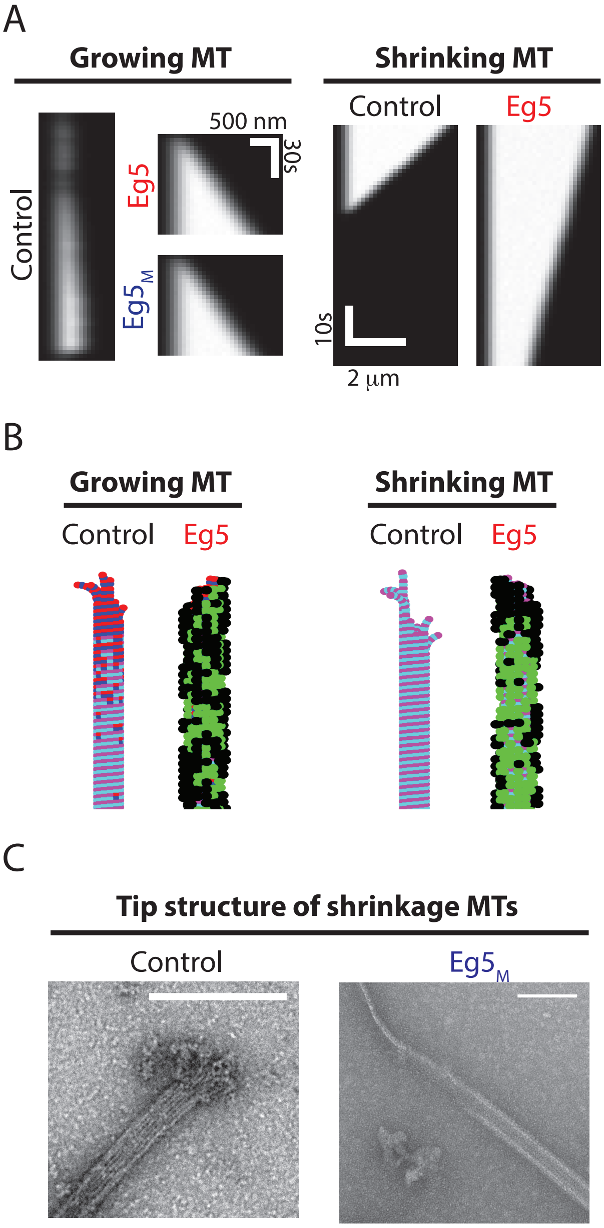


**Figure S5. Related to Figure 5. Parameterization of simulated microtubule dynamics.** (A) Simulated kymographs of growing and shrinking microtubules. (B) Illustration of Eg5-stimulated microtubule stability. Microtubule tip contains sheet structure in the presence of Eg5, whereas generates ram's horn in control (Red and blue pair: GTP-tubulin; magenta and cyan pair: GDP-tubulin; green: two-heads-bound Eg5; black: one-head-bound Eg5). (C) EM images of shrinking microtubules in the presence or absence of monomeric Eg5. The diverse tips in the presence or absence of motors are consistent with the model illustration in **Figure S5C**. Scale bar: 100 nm.

**Table S1. Parameters for Eg5-stimulated microtubule polymerization model.**

| Rate constants | Description | Source | Value | Unit |
|---|---|---|---|---|
| **Parameters for microtubule dynamics** | | | | |
| $k_{on}$ | Longitudinal tubulin-tubulin on-rate | Constrained¶ | 1.95 | $\mu M^{-1}s^{-1}$ per MT |
| $k_{off}^{GTP}$ | Longitudinal tubulin off-rate in GTP | Constrained¶ | 3 | $s^{-1}$ |
| $k_{off}^{GDP}$ | Longitudinal tubulin off-rate in GDP | Constrained¶ | 5 | $s^{-1}$ |
| $k_{zip}$ | Lateral bond annealing rate | Constrained¶ | 10 | $s^{-1}$ |
| $k_{unzip}$ | Lateral bond breaking rate | Constrained¶ | 30 | $s^{-1}$ |
| $k_{hyd}$ | Tubulin GTP hydrolysis rate | Melki et al., 1996 | 0.05 | $s^{-1}$ |
| $F_{GTP}$ | GTP-induced lateral bond stabilization factor | Constrained¶ | 2.5 | fold |
| $F_{wall}$ | Mechanical stabilization by Mt lattice | Margolin et al., 2012 | 9 | fold |
| **Parameters for dimeric Eg5-microtubule interaction** | | | | |
| $k_{on}^{Motor}$ | Eg5-microtubule binding rate | Chen et al. 2016 | 20 | $\mu M\ MT^{-1}s^{-1}$ |
| $k_{step}$ | Eg5 stepping rate | Chen et al. 2016 | 10 | $s^{-1}$ |
| $k_{unbind}$ | Eg5-microtubule unbinding rate | Chen and Hancock 2015 | 0.25 | $s^{-1}$ |
| $k_{slow}$ | Eg5 stepping rate on curved protofilaments | Constrained by **Figure 3G** | 1 | $s^{-1}$ |
| $k_{pause}$ | Eg5 unbinding rate on curved protofilaments | Chen and Hancock 2015 | 0.1 | $s^{-1}$ |
| **Parameters for monomeric Eg5-microtubule interaction** | | | | |
| $k_{on}^{Motor}$ | $Eg5_M$-microtubule binding rate | Chen et al. 2016 | 10 | $\mu M\ MT^{-1}s^{-1}$ |
| $k_{unbind}$ | $Eg5_M$-microtubule unbinding rate | Chen et al. 2016 | 10 | $s^{-1}$ |
| $k_{pause}$ | $Eg5_M$ unbinding rate on curved protofilaments | Chen and Hancock 2015 | 0.5 | $s^{-1}$ |
| **Parameters for Eg5-induced microtubule polymerization** | | | | |
| $F_{zip}^{Eg5}$ | Eg5 enhancement of lateral bond formation | Constrained by **Figure 2E** | 2.2 | fold |
| $F_{unzip}^{Eg5}$ | Eg5 slowing of lateral bond breakage | Constrained by **Figure 2F** | 10 | fold |

¶: Parameters for simulating microtubule dynamics were trained by MT growth assays (**Figure 2E**), GDP-MT shrinkage assays (**Figure 2F**), GMPCPP-MT shrinkage rate (Chen and Hancock, 2015), and fray size in both growth and shrinkage microtubules (McIntosh et al., 2018). In addition to setting lateral bond zipping rate for GDP-tubulin and GDP-lattice as zero, no additional degrees of freedom are required.

**Table S2. Kinetics and free energy of tubulin lattice.**

| Description | | Association rate ($k_{on}$) | Dissociation rate ($k_{off}$) | Dissociation constant ($K_D$) | Free energy ($\Delta G^*$) |
|---|---|---|---|---|---|
| **Thermodynamics of microtubule dynamics** | | | | | |
| Longitudinal | GTP | 0.15 ($\mu M^{-1}s^{-1}$) | 3 ($s^{-1}$) | $2 \times 10^{-6}$ M | -13.1 $k_BT$ |
| | GDP | 0.15 ($\mu M^{-1}s^{-1}$) | 5 ($s^{-1}$) | $3.3 \times 10^{-6}$ M | -12.6 $k_BT$ |
| Lateral | GDP\|GDP | 0 | 30 ($s^{-1}$) | NA | NA |
| | GDP\|GTP¶ | 10 ($s^{-1}$) | 12 ($s^{-1}$) | 1.2 | 0.2 $k_BT$ |
| | GTP\|GTP | 10 ($s^{-1}$) | 4.8 ($s^{-1}$) | 0.48 | -0.7 $k_BT$ |
| Lateral, wall§ | GDP\|GDP | 0 | 3.3 ($s^{-1}$) | NA | NA |
| | GDP\|GTP¶ | 10 ($s^{-1}$) | 1.3 ($s^{-1}$) | 0.13 | -2.0 $k_BT$ |
| | GTP\|GTP | 10 ($s^{-1}$) | 0.5 ($s^{-1}$) | 0.05 | -3.0 $k_BT$ |
| Contribution of each Eg5 motor domain | | | | | -3.1 $k_BT$ |

¶: Ignores the influence of microtubule chirality.

§: Wall denotes the lateral zipping process between the two sheets or bundles.
